# Defects in Mitochondrial Biogenesis Drive Mitochondrial Alterations in PINK1-deficient Human Dopamine Neurons

**DOI:** 10.1101/2023.06.23.546087

**Authors:** Hu Wang, Rong Chen, Liming Xiao, Manoj Kumar, Jesús Acevedo-Cintrón, Joanna Siuda, Dariusz Koziorowski, Zbigniew K. Wszolek, Valina L. Dawson, Ted M. Dawson

## Abstract

Mutations and loss of activity in the protein kinase PINK1 play a role in the pathogenesis of Parkinson’s disease (PD). PINK1 regulates many aspects of mitochondrial quality control including mitochondrial autophagy (mitophagy), fission, fusion, transport, and biogenesis. Defects in mitophagy are though to play a predominant role in the loss of dopamine (DA) neurons in PD. Here we show that, although there are defects in mitophagy in human DA neurons lacking PINK1, mitochondrial deficits induced by the absence of PINK1 are primarily due to defects in mitochondrial biogenesis. Upregulation of PARIS and the subsequent down regulation of PGC-1α accounts for the mitochondrial biogenesis defects. CRISPR/Cas9 knockdown of PARIS completely restores the mitochondrial biogenesis defects and mitochondrial function without impacting the deficits in mitophagy due to the absence of PINK1. These results highlight the importance mitochondrial biogenesis in the pathogenesis of PD due to inactivation or loss of PINK1 in human DA neurons.

## Introduction

Parkinson’s disease (PD) is the second most prevalent neurological disease caused by progressive loss of dopaminergic (DA) neurons in the substantial nigra [1]. Mutations in several genes encoding PINK1 (PTEN-induced novel kinase 1), LRRK2 (leucine rich repeat kinase 2), parkin, DJ-1, and α-synuclein have been associated with autosomal dominant familial PD [2, 3]. However, the exact mechanism underlying cell death of DA neurons in PD remain elusive.

Recent studies suggest that the PINK1/parkin pathway is implicated in mitochondrial quality control [4–6]. By using the immortal human HeLa cancer cell line, PINK1 was identified to recruit parkin to the outer membrane of damaged mitochondria in response to depolarizing toxins [7, 8]. These initial investigations offered new insight of the underlying mechanisms of PINK1/parkin in maintaining mitochondrial health through promoting ubiquitin-dependent mitochondrial autophagy (mitophagy) [9, 10]. Loss-of-function mutations in PINK1 and parkin failed to mediate mitophagy in cultured cells and neurons [11, 12]. Thus, the hypothesis of defective mitophagy has been a leading theory in PINK1 or parkin deficient DA neuron neurodegeneration associated with PD.

PINK1 and parkin work in a coordinated manner in mitophagy in ex vivo culture systems, however, whether PINK1-PARKIN-directed mitophagy occurring in vivo, and especially whether progressive DA neuron loss is primarily due to mitophagy defects is not known [11]. Growing evidence indicates that the accumulation of parkin interacting substrate (PARIS, ZNF746) drives the progressive loss of DA neurons in the ventral midbrain of adult mice [13–17], in Drosophila [18] lacking PINK1 or PARKIN, and in A53T α-synuclein (α-syn) transgenic models of familial PD as well in the α-syn preformed fibril (PFF) sporadic PD model [14, 19, 20] through downregulation of the peroxisome proliferator-activated receptor gamma, coactivator 1α (PGC-1α) which was recognized as a “master regulator” of mitochondrial biogenesis [21]. In addition mitochondrial biogenesis seems to drive DA neuron loss and dysfunction due to parkin deficiency in human DA neurons in the pathogenesis of PD [22]. Whether similar processes occur in human DA neurons lacking PINK1 is not known. Here, using human DA neurons lacking PINK1, we explored the role of defects in mitochondrial biogenesis versus mitophagy and find that defects in mitochondrial biogenesis is the major driver of mitochondrial dysfunction and human DA neuron dysfunction due to inactivation or loss of PINK1.

## Results

### Human DA neurons lacking PINK1 have reduced levels of DA markers compared to isogenic Control

PINK1 deficient (I368N 1#) and its isogenic control (iControl) inducible pluripotent stem cells (iPSCs) lines [23] and two additional human PINK1 deficient iPSC (PINK1 I368N 1# and Q456X [24]) and two human iPSC control lines (SC1014 and SC1015) were differentiated into neural progenitor cells (NPC) followed by differentiation into DA neurons (Figure S1A and S1B) [22]. Administration of FCCP increases the expression of PINK1 leading to increased p-Ser65-Ub and reduced parkin levels (Figure S1D-F) in iControl DA neurons, while PINK1 levels were not substantially increased nor was there an elevation in p-Ser65-Ub in the PINK1 deficient (I368N 1#) neurons (Figure S1C-F). In the human SC1014 line, FCCP increased both PINK1 and p-Ser65-Ub levels and levels of parkin were decreased (Figure S1G-J), while in the PINK1 Q456X deficient line administration of FCCP failed to increase the expression of PINK1 and there was no increase in p-Ser65-Ub or change in parkin levels (Figure S1G-J). Similar results were obtained in the human SC105 control line and the PINK1 I368N #2 line respectively (Figure S1G-J). There was a significant 51.72% reduction of the percentage of tyrosine hydroxylase (TH) positive neurons in the PINK1 I368N1# deficient line compared to its iControl line (Figure 1A, B), while there was only a 14.61% reduction in overall neuronal number as assessed by TUJ1 immunoreactivity (Figure 1C). Of the TUJ1 positive cells there is a 50.75% reduction in the number of TH positive neurons in the PINK1 I368N 1# deficient neuronal cultures compared to the isogenic control neuronal cultures (Figure 1D). Immunoblot analysis confirmed the loss of TH, TUJ1 and relative TH/TUJ1 levels (Figure 1E to H). To determine whether the loss of TH was due to cell death staining with propidium iodide (PI), a marker of cell death was performed [25]. In the PINK1 deficient I368N 1# line compared to the iControl line there was an increase of PI staining (19.9%) indicative of cell death (Figure 1I, J).

**Figure 1.**
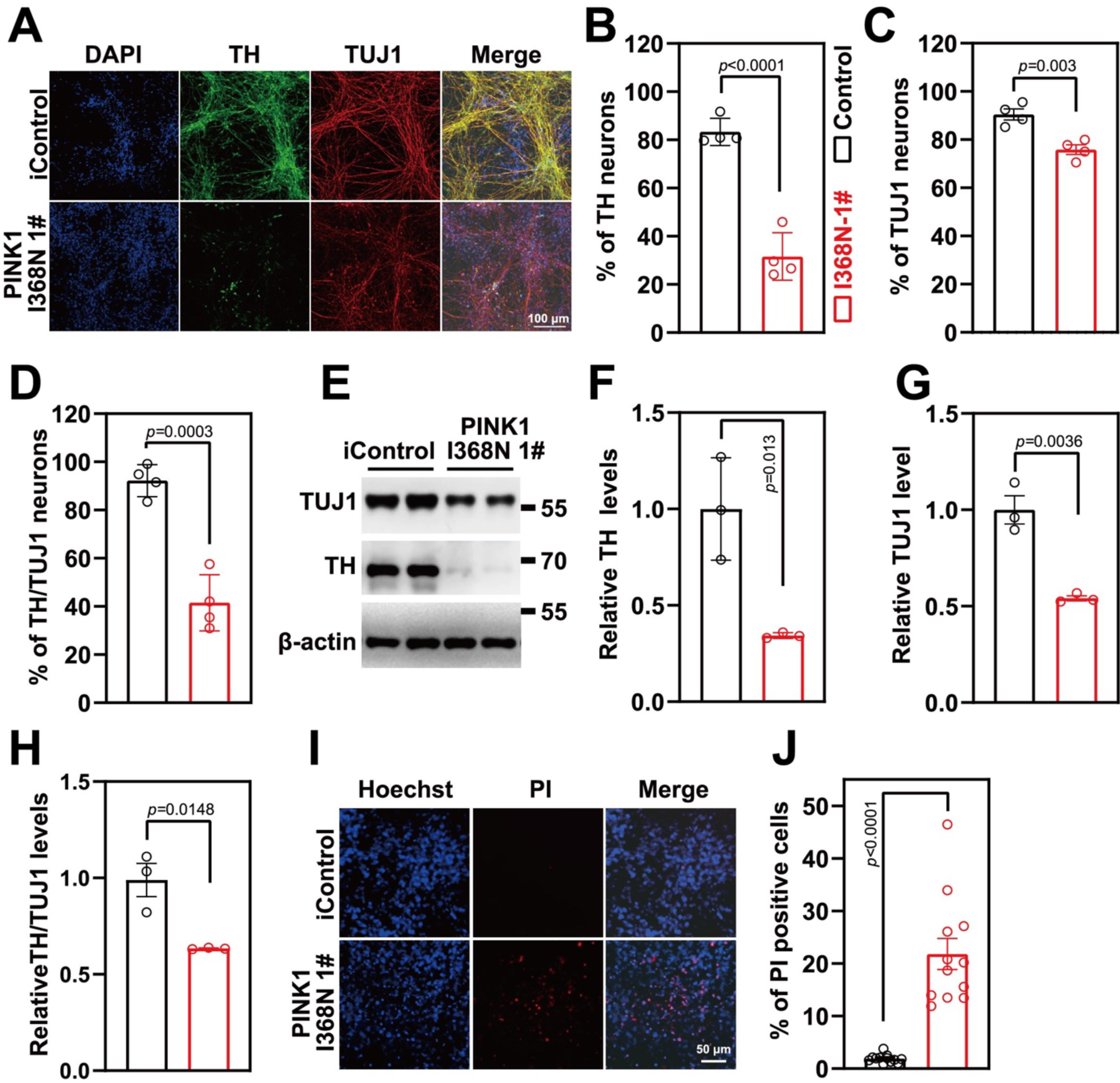
DA neuron differentiation and characterization. A. Representative images of differentiated DA neuron at day 60 from isogenic control and PINK1 p.I368N 1# mutant lines for neurons (TUJ1, Red) and dopaminergic neurons (TH, green), DAPI staining in blue represents the nucleus. B. Quantification of TH positive neuron percentage of immunostaining. C. Quantification of TUJ1 positive neuron percentage of immunostaining. D. Quantification of TH/TUJ1 double positive neuron percentage of immunostaining. E. Immunoblot analysis of TH and TUJ1 protein level in differentiated DA neurons. F. Quantified intensities of TH are shown, relative protein level was normalized to β-actin. N=3 independent experiments. G. Quantified intensities of TUJ1 are shown, relative protein level was normalized to β-actin. N=3 independent experiments. H. Quantified intensities of TH/TUJ1 are shown. I. Representative images of PI staining, DAPI is in blue, PI is in red. J. Quantification of PI staining. N=3 independent experiments.

### PINK1 deficiency leads to mitochondrial respiratory decline

Mitochondrial respiration was measured in the human DA neuron cultures using a Seahorse XF24 Flux Analyzer (Agilent Technologies, USA). There was a 13.59% reduction in basal respiration, a 30% reduction in spare respiratory capacity and a 46.32% reduction in carbonyl cyanide m-chlorophenyl hydrazine (FCCP)-induced maximal respiration in the PINK1 I368N 1# deficient neuronal cultures compared to the iControl culture (Figure 2A and 2B). In addition, there was a 26.73% reduction in basal respiration, an 80% reduction in spare respiratory capacity and a 75.89% reduction in FCCP-induced maximal respiration in the PINK1 Q456X deficient iPSC derived DA neurons compared to the iPSC SC1014 control DA neurons (Figure S2A-S2C). Similar results were obtained in the human SC105 control line and the PINK1 I368N #2 line respectively (Figure S2D-F). Mitochondrial membrane potential assessment using MitoCMXRos revealed a 91.6% reduction in the PINK1 I368N 1# culture compared to the iControl culture (Figure 2D and 2E). Similarly, a 75.2% reduction in membrane potential was found in the PINK1 Q456X deficient iPSC derived DA neurons compared to the iPSC SC1014 control DA neurons (Figure S2G-S2H). Similar results were obtained in the human SC105 control line and the PINK1 I368N 1# line respectively (Figure S2I-J). Taken together these results indicate there is a deficit in mitochondrial respiration in PINK1 deficient DA neurons.

**Figure 2.**
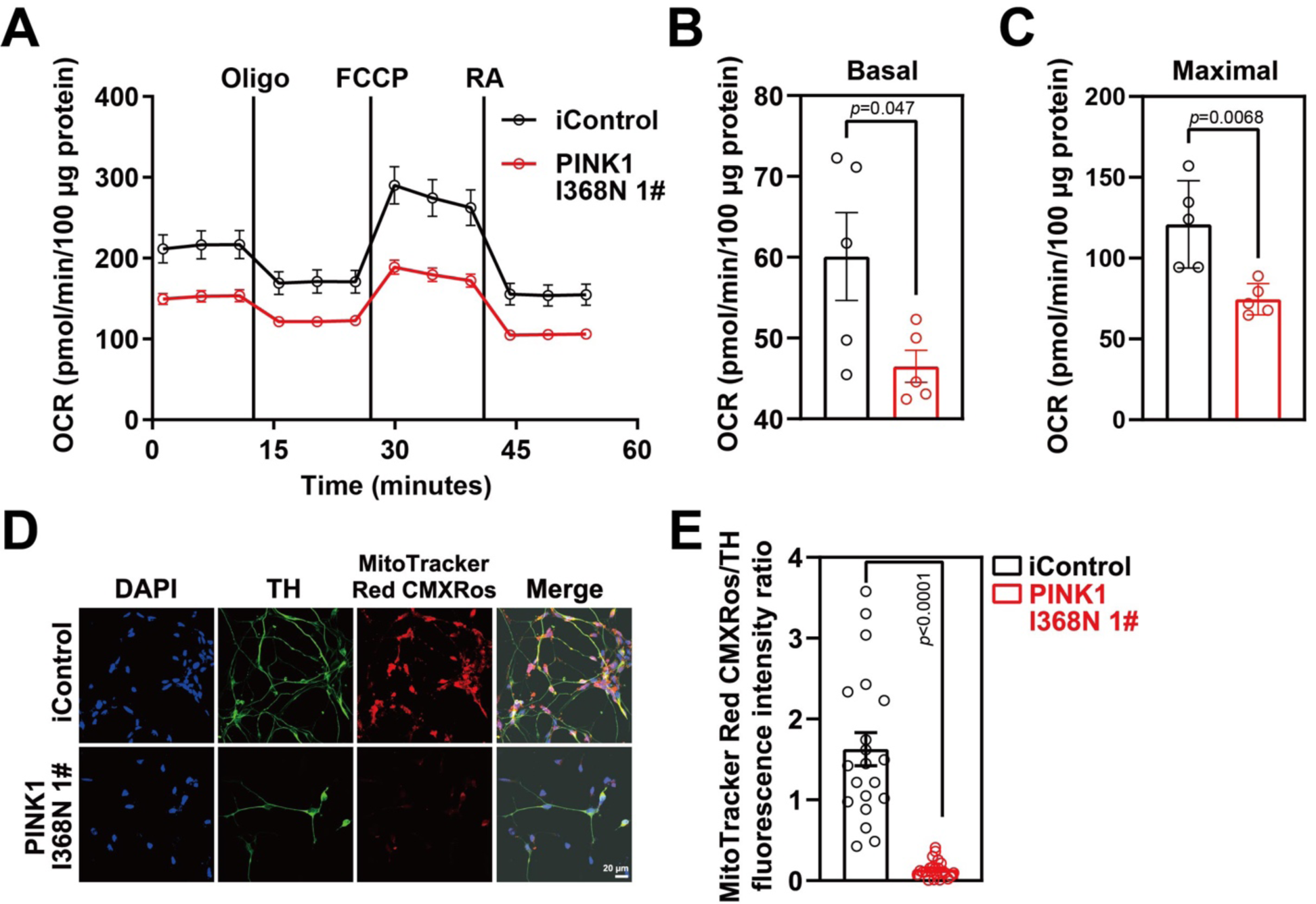
Mitochondrial dysfunction in PINK1 p.I368N-1# mutant DA neurons. A. Mitochondrial oxygen consumption rate curve generated using Seahorse platform showing the mitochondrial dysfunction in isogenic control and PINK1 p.I368N 1# mutant neurons in the presence of oligomycin, FCCP, and rotenone respectively. B. Quantification of basal respiration in isogenic control and PINK1 p.I368N-1# mutant neurons. N=3 independent experiments. C. Quantification of maximal respiration in isogenic control and PINK1 p.I368N 1# mutant neurons. N=3 independent experiments. D. Representative immunostaining images of TH-positive neurons and MitoTracker Red CMXRos from isogenic control and PINK1 p.I368N 1# mutant neurons. E. Quantification of TH and MitoTracker Red CMXRos intensity. N=20 TH positive neurons in each group. N=15-30 TH positive neurons in each group.

### Reduced mitochondrial autophagy in PINK1 deficient human DA neuron culture

Mitophagy was measured in human DA neuron cultures by utilizing Mito-Keima [26]. Human DA neuron cultures were transduced with a lenti virus expressing Mito-Keima, a pH sensitive marker of mitophagy that provides a measure of the status of mitochondria where mitochondria that are localized to cytoplasm are green while those mitochondria that are being degraded via autophagy are red due to the acidic environment of the lysosome [26]. The ratio of the lysosomal red signal over the mitochondrial green signal within the neuronal body was calculated as the mitophagy index. There was a 78.4% reduction in the mitophagy index in the PINK1 I368N 1# deficient DA neurons compared to the iControl DA neurons (Figure 3A and 3B). There was an 86.8% reduction in the mitophagy index of the PINK1 Q456X deficient iPSC derived DA neurons compared to the iPSC SC1014 control DA neurons (Figure S3A-S3B). Similar results were obtained in the human SC105 control line and the PINK1 I368N 1# line, respectively (Figure S3C-S3D). We evaluated LC3I/II levels, an integral structural protein of the autophagasome for autophagy. The LC II ratio to LCI is decreased by 85.4% in the PINK1 I368N 1# deficient DA neurons compared to the iControl DA neurons (Figure 3C and 3D) and the levels of the p62 protein significantly accumulated in PINK1 I368N 1# deficient neurons compared to the iControl DA neurons (Figure 3C and 3E). There was a 58.4% reduction in the LC II ratio to LCI and an elevation of p62 levels in the PINK1 Q456X and PINK1 I368N 1# deficient iPSC derived DA neurons compared to the iPSC SC1014 and SC1015 control neurons (Figure S3E-S3B). These results are consistent with the absence of PINK1 leading to mitophagy defects.

**Figure 3.**
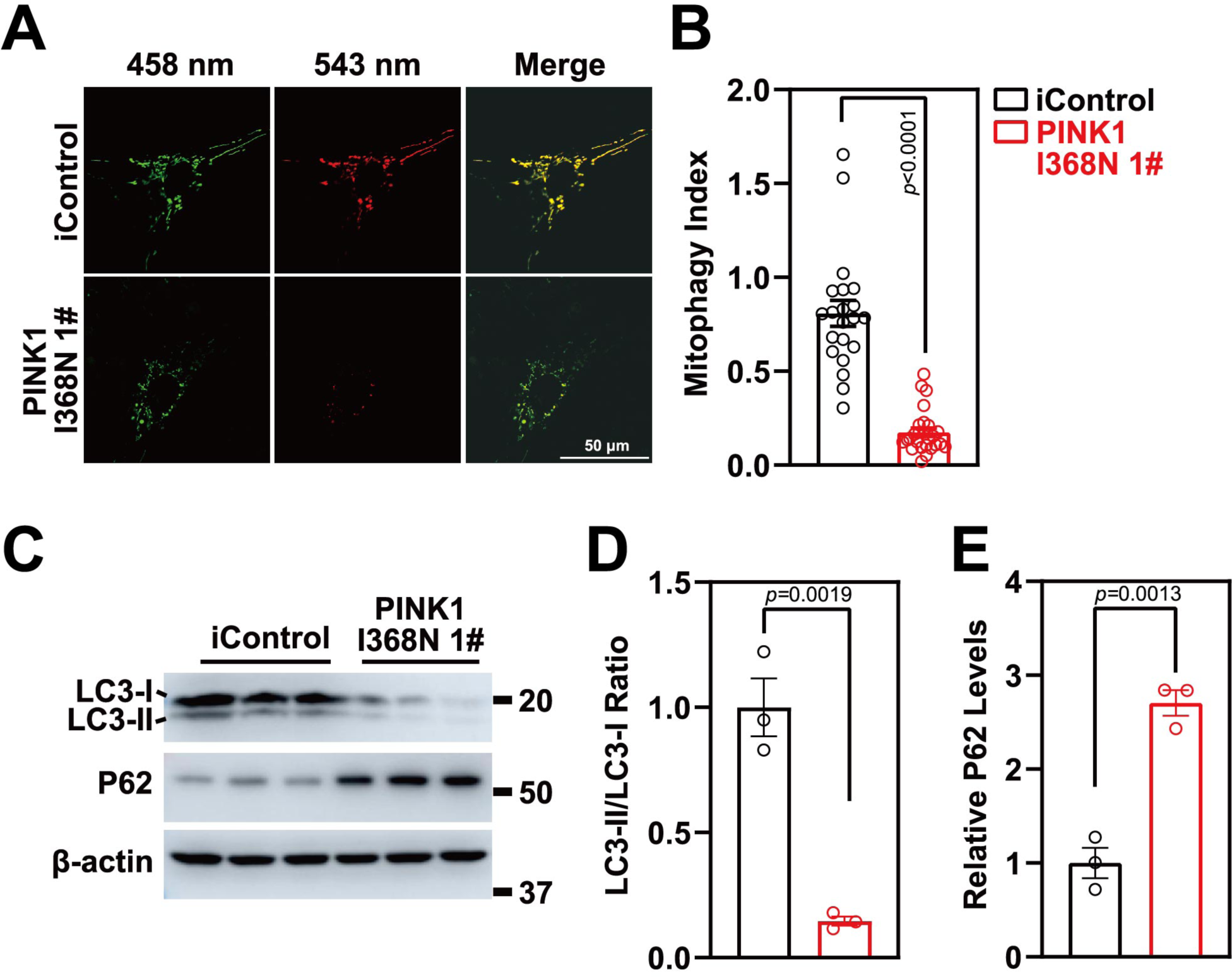
Mitophagy defects in PINK1 p.I368N-1# mutant neurons. A. Representative confocal live images of isogenic control and PINK1 p.I368N 1# mutant neurons infected with lentivirus encoding for MitoKeima. B. Quantified mitophagy index in isogenic control and PINK1 p.I368N-1# mutant neurons. N=20 neurons in each group. C. Analysis of LC3-I/II and autophagy marker P62 protein level by immunoblot. A. Quantification of immunoblot of LC3-I/II ratio are shown. N=3 independent experiments. B. Quantification of immunoblot of P62 normalized to β-actin. N=3 independent experiments.

### Elevation of PARIS and Reduction of PGC-1α in PINK1 deficient human DA neuron cultures

The level of PARIS was elevated 2-fold in the PINK1 I368N 1# deficient DA neurons compared to the iControl DA neuronal cultures (Figure 4A and 4B). PARIS is a transcriptional repressor that regulates the expression of peroxisome proliferator-activated receptor gamma, coactivator 1α, PGC-1α, a master coregulator of mitochondrial biogenesis, oxidative stress management and function [21]. There was an 81.73% reduction in PGC-1α levels in PINK1 I368N 1# deficient DA cultures compared to the iControl neuronal cultures (Figure 4A and 4C). Accompanying the elevation in PARIS and reduction in PGC-1α was a signification 59.35% reduction in TH expression via immunoblot analysis (Figure 4A and D). In addition, there is a 2-fold increase in PARIS levels in the Q456X and PINK1 I368N 1# deficient iPSC derived DA neuron cultures compared to the iPSC SC1014 and SC1015 control neurons (Figure S4A and S4B). In the PINK1 Q456X and PINK1 I368N 1# deficient iPSC derived DA neuron cultures there is significant reduction in TH immunoreactivity compared to the iPSC SC1014 and SC1015 control DA neurons (Figure S4A and S4B). There is a 72.1% reduction in PGC-1α levels in the PINK1 Q456X and PINK1 I368N 1# deficient iPSC derived DA neuron cultures compared to the iPSC SC1014 and SC1015 control DA neuron cultures (Figure S4A and S4B). Transmission electron microscopy (TEM) of mitochondria from PINK1 I368N 1# deficient DA neurons compared to the iControl human DA neuronal cultures reveals that the mitochondria are significantly reduced in size and number (Figure 4E to 4G). Mitochondrial copy number was assessed by measuring the mitochondrial genes, tRNA Leu (MT), NADH-ubiquinone oxidoreductase chain 1 (ND1), mitochondrially encoded ATP synthase membrane subunit 6 (ATP6) and mitochondrially encoded cytochrome c oxidase I (COX1), normalized to nuclear encoded β2-microglobulin gene. There is a 50% reduction in tRNA Leu (MT), ND1, ATP6 and COX1 levels in the PINK1 deficient neuron compared to the iControl neuronal cultures (Figure 4H). There was also a significant reduction in the integral mitochondrial proteins, cytochrome C (Cyt C) and CoxIV levels in the PINK1 I368N 1# deficient DA neurons compared to the iControl human DA neuronal cultures (Figure 4I and 4J). There was a 49% reduction in MT, ND1, ATP6 and COX1 levels in the PINK1 Q456X deficient iPSC derived DA neurons compared to the iPSC SC1014 control DA neurons (Figure S4C). Similar results were obtained in the human PINK1 I368N 1# line and SC105 control line, respectively (Figure S4D). Taken together these results indicate that there is a reduction in mitochondrial mass in PINK1 deficient human DA neurons.

**Figure 4.**
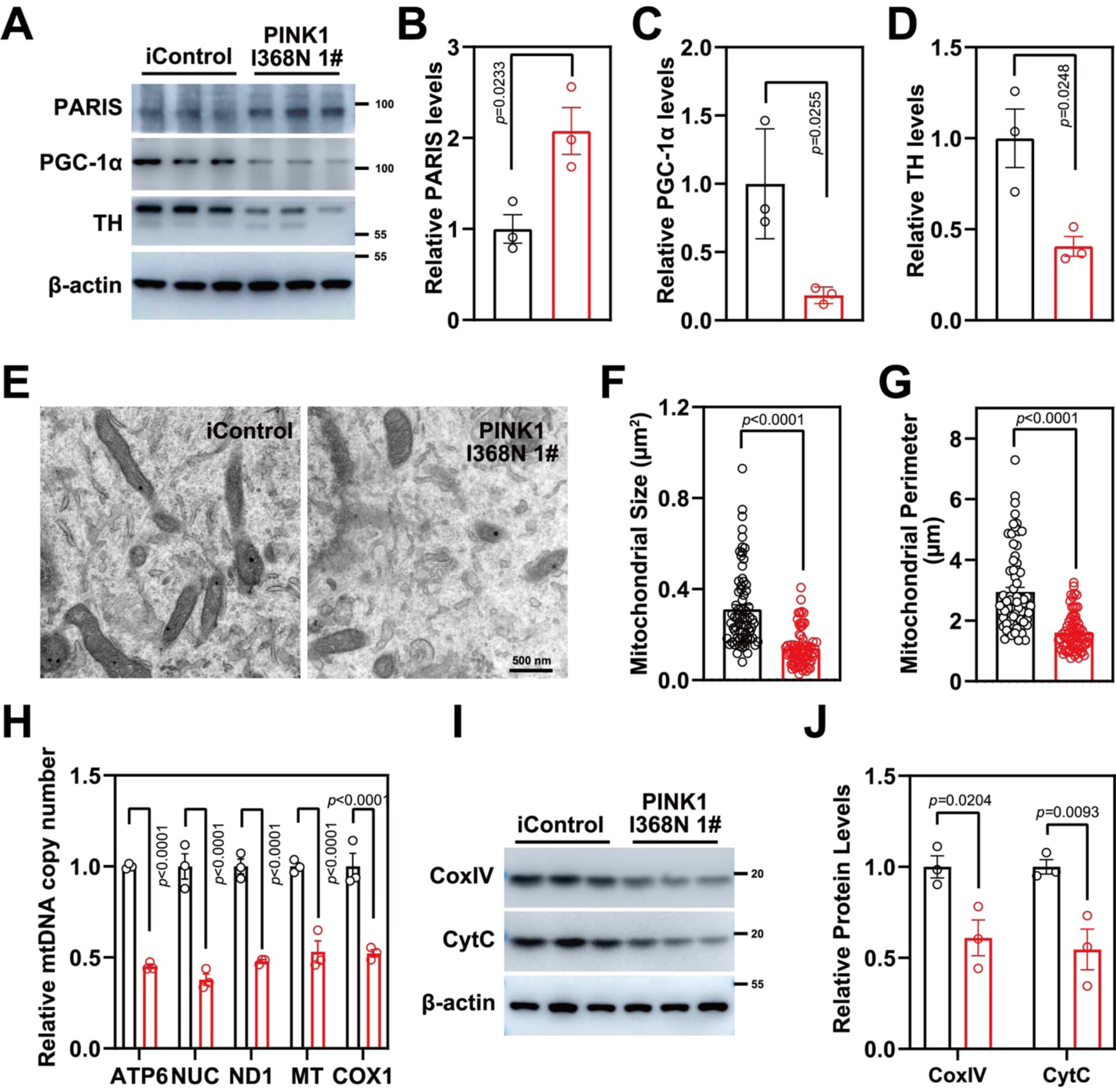
Mitochondrial biogenesis defects in PINK1 p.I368N-1# mutant neurons. A. Analysis of PARIS, PGC-1α and TH protein level in isogenic control and PINK1 p.I368N 1# mutant neurons by immunoblot. C. Quantification of immunoblot of PARIS normalized to β-actin. N=3 independent experiments. D. Quantification of immunoblot of PGC-1α normalized to β-actin. N=3 independent experiments. E. Quantification of immunoblot of TH normalized to β-actin. N=3 independent experiments. B. Representative transmission electron microscopy images of neurons from isogenic control and PINK1 p.I368N 1# mutant. C. Quantification of mitochondrial size per cell body in isogenic control and PINK1 p.I368N 1# mutant neurons. N=15-30 TH positive neurons in each group. D. Quantification of mitochondrial perimeter per cell body in isogenic control and PINK1 p.I368N 1# mutant neurons. N=15-30 TH positive neurons in each group. F. Relative mitochondrial DNA copy number quantification of ATP6, NUC, ND1, MT, and COX1 in isogenic control and PINK1 p.I368N 1# mutant neurons. N=3 independent experiments. E. Analysis of CoxIV and CytC protein level in isogenic control and PINK1 p.I368N 1# mutant neurons by immunoblot. G. Quantification of immunoblot of CoxIV and CytC in isogenic control and PINK1 p.I368N 1# mutant neurons. N=3 independent experiments.

### Mitochondrial biogenesis defect in PINK1 deficient human DA neurons

A (SNAP-TAG) self-protein labeling tag covalently fused with suitable ligand targeted to the mitochondria via fusion to the integral mitochondrial protein Cox8A was transduced into human DA neuron cultures via lentivirus to monitor mitochondrial turnover [27, 28]. Old versus new mitochondria were identified by using a red (TMR-Star) SNAP-TAG and a green (Oregon Green) SNAP-Tag substrate, in TH positive neurons and demonstrates that there was a significant 76.7% reduction in the formation of new mitochondria within PINK1 I368N 1# deficient DA neurons compared to the iControl DA neuronal cultures (Figure 5A and 5B). A modified non-radioactive protein puromycin labelling called surface sensing of translation (SUnSET) assay [29] was performed by assessing TOM20 and puromycin immunofluorescence levels within TH positive neurons was also used to assess mitochondrial biogenesis in isogenic control and PINK1 p.I368N 1# mutant neurons (Figure 5C and 5D). There is a 52.4% reduction of puromycin labeling in TOM20 labeled mitochondria in human DA neurons (Figure 5C and 5D). There was significant 86.9% reduction in the formation of new mitochondria within DA neurons in the PINK1 Q456X and PINK1 I368N 1# deficient iPSC derived DA neuron cultures compared to the iPSC SC1014 and SC1015 control DA neuron cultures (Figure S4E to S4G), and 56.2% reduction of puromycin labeling in Tom20 labelled mitochondria in PINK1 deficient DA neurons (Figure S4H to S4J). However, we observed that there was no significant defect in mitochondrial biogenesis in non-TH positive neurons examined by SNAP-TAG assays (Figure S4K and Figure S4L) and SUnSET (Figure S4K and Figure S4M).

**Figure 5.**
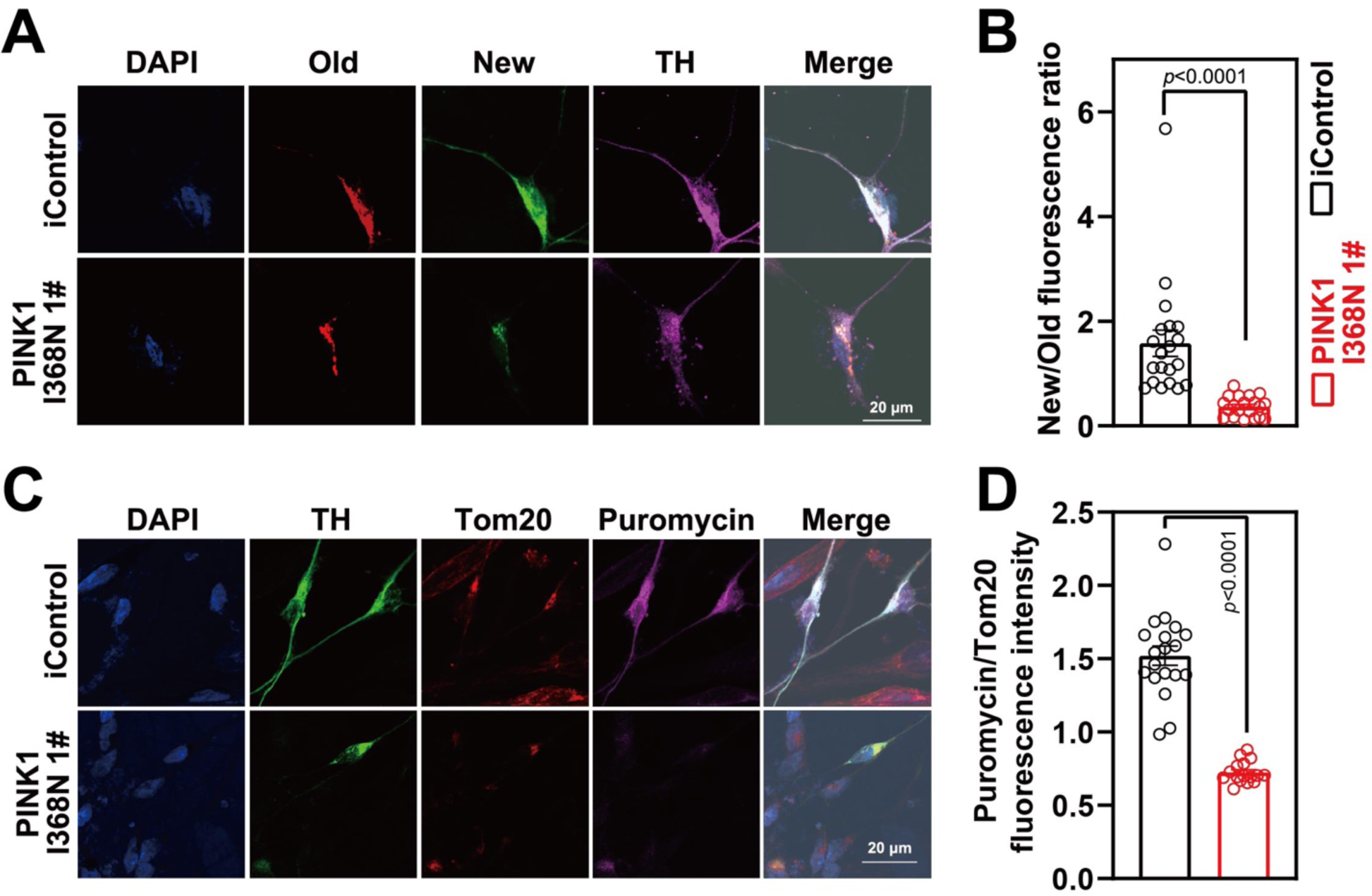
Defects of mitochondrial biogenesis in PINK1 p.I368N-1# mutant DA neurons. A. Representative confocal images of SNAP-TAG Cox8a labeled mitochondria with TH staining in isogenic control and PINK1 p.I368N 1# mutant neurons. B. Fluorescence intensity quantification of old and new mitochondria labeled by Cox8a in TH positive neurons. N=20 TH positive neurons in each group. C. Representative confocal images of puromycin labelling SUnSET assay in isogenic control and PINK1 p.I368N 1# mutant neurons. D. Immunofluorescence intensity quantification of Tom20 and puromycin within TH positive neurons. N=20 TH positive neurons in each group.

### Reducing PARIS levels restores PGC-1α levels without restoring the defects in mitophagy

PARIS levels were reduced in human DA neuron cultures via CRISPR/Cas9 using lentiviral transduction of a guide RNA to PARIS. The levels of PARIS were reduced by 77% in PINK1 I368N 1# deficient DA neurons compared to PINK1 I368N 1# deficient DA neurons without the PARIS guide RNA (Figure 6A and 6B). There was also a 61.7% reduction in the PINK1 Q456X and PINK1 I368N 1# deficient iPSC derived DA neuron / PARIS KD iPSC derived DA neurons compared to the PINK1 Q456X and PINK1 I368N 1# deficient iPSC derived DA neurons (Figure S5A to S5B). Accompanying the reduction in PARIS levels in the PINK1 I368N 1# deficient DA transduced with a guide RNA to PARIS was a restoration of PGC-1α and TH levels in the PINK1 I368N 1# deficient DA neurons compared to PINK1 I368N 1# deficient DA neurons without the PARIS guide RNA (Figure 6A, 6C and 6D). A similar restoration of PGC-1α levels were observed in the PINK1 Q456X and PINK1 I368N 1# deficient iPSC derived DA neurons transduced with the guide RNA to PARIS (Figure S5A and S5C). The mitophagy index remained significantly reduced in the PINK1 I368N 1# deficient in the setting of PARIS knock down (Figure 6G and 6H). Levels of p62 and LC3 I/II levels also remain unchanged following PARIS knock down (Figure 61, 6J and 6J). The mitophagy index was not reduced by PARIS knock down in PINK1 Q456X and PINK1 I368N 1# deficient iPSC derived DA neurons Figure S5D-S5G, while PINK1 overexpression rescues the mitophagy index (Figure S5H to S5L).

**Figure 6.**
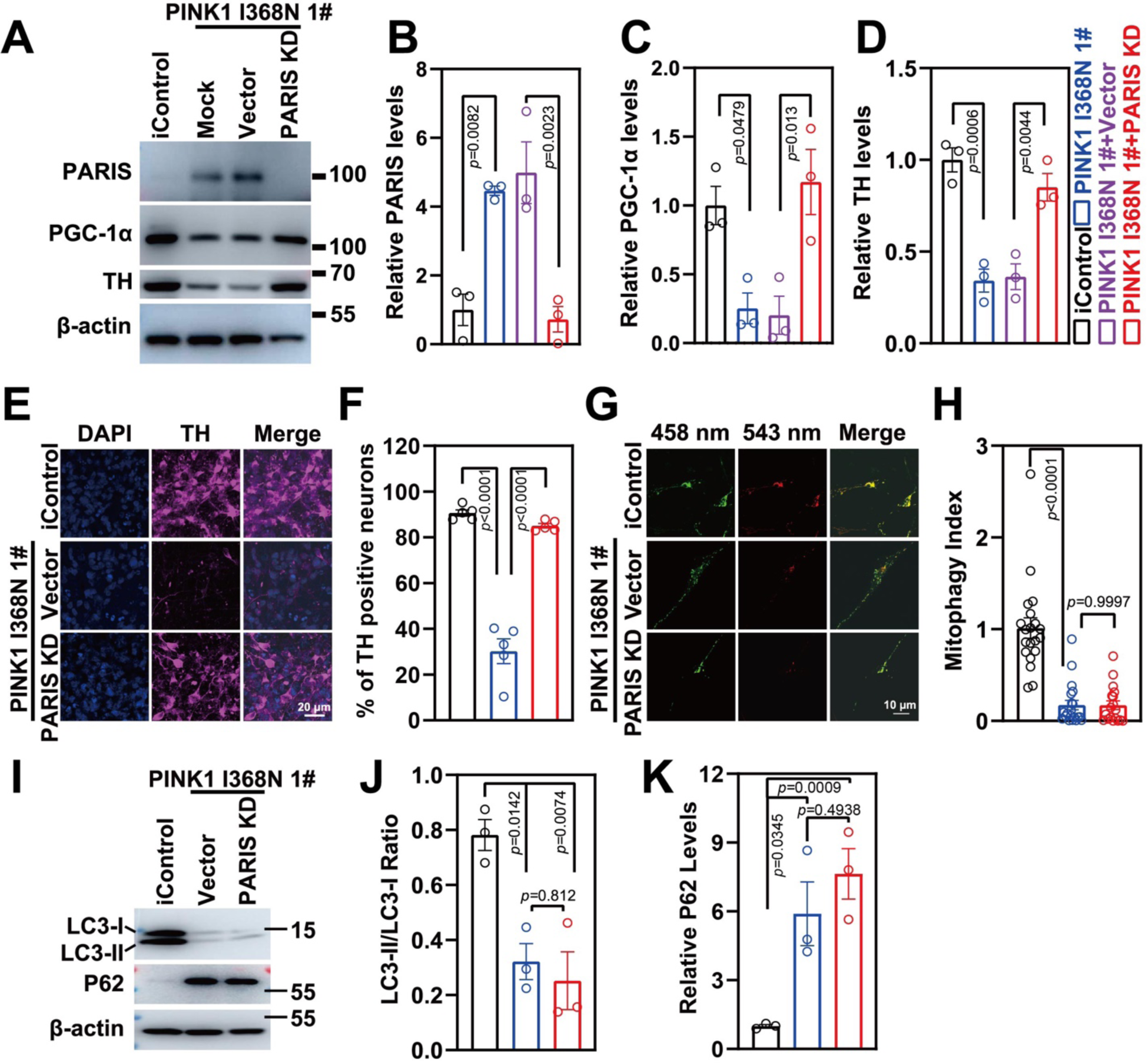
Characterization of PARIS knockdown in PINK1 p.I368N-1# mutant neurons. H. Immunoblot analysis of PARIS, PGC-1α and TH protein level in isogenic control and PINK1 p.I368N 1# mutant neurons with or without PARIS knockdown. I. Quantified intensities of PARIS are shown, relative protein level was normalized to β-actin. N=3 independent experiments. J. Quantified intensities of PGC-1α are shown, relative protein level was normalized to β-actin. N=3 independent experiments. K. Quantified intensities of TH are shown, relative protein level was normalized to β-actin. N=3 independent experiments. L. Representative confocal images of anti-TH immunostaining in isogenic control and PINK1 p.I368N 1# mutant neurons. A. Quantification of TH positive neurons by anti-TH immunostaining in isogenic control and PINK1 p.I368N 1# mutant neurons. N=20 TH positive neurons in each group. M. Representative confocal images of isogenic control and PINK1 p.I368N 1# mutant neurons infected with lentivirus encoding Mito-Keima mitophagy reporter systems. B. Quantified mitophagy index in isogenic control and PINK1 p.I368N 1# mutant neurons with or without PARIS knockdown. N=20 TH positive neurons in each group. N. Immunoblot analysis of LC3-I/II and P62 protein level in isogenic control and PINK1 p.I368N 1# mutant neurons with or without PARIS knockdown. O. Quantification of LC3-I/II and P62 protein levels. N=3 independent experiments.

### Reducing PARIS levels restores the mitochondrial respiratory and biogenesis defects in human DA neuron deficient in PINK1

The reduction in basal respiration, spare respiratory capacity and maximal respiration is rescued in the PINK1 I368N 1# deficient DA neurons by knocking down PARIS (Figure 7A-C). Basal respiration, spare respiratory capacity and maximal respiration is also rescued in the in PINK1 Q456X and PINK1 I368N 1# deficient iPSC derived DA neuron cultures (Figure S6A-F). Knockdown of PARIS restores the mitochondrial biogenesis defect in DA neurons as determined by the puromycin labelling SUnSET (Figure 7D and 7E) and SNAP-tag assays in the PINK1 I368N 1# deficient DA neurons (Figure 7F and 7G) as well as in the Q456X and PINK1 I368N 1# deficient iPSC derived DA neuron cultures (Figure S7A to S7H). Mitochondrial copy number as assessed by MT, ND1, ATP6 and COX1 levels normalized to nuclear encoded β2-microglobulin gene is restored in the PINK1 I368N 1# deficient DA neurons (Figure S7G) and in the PINK1 Q456X and PINK1 I368N 1# deficient iPSC derived DA neuron cultures (Figure S7I and S7J). The reduction in size and number as well as the reduction in the percentage of healthy mitochondria was significantly restored by PARIS knockdown in the PINK1 I368N 1# DA neurons (Figure 7I, 7J and 7K). The reduction in levels of the outer mitochondrial protein TOM20 in TH positive neurons is significantly restored by PARIS knockdown in PINK1 I368N 1# deficient DA neurons (Figure 7L and 7M). Reduction in mitochondrial membrane potential in PINK1 Q456X and PINK1 I368N 1# deficient iPSC derived DA neurons and rescued by PARIS KD (Figure S6G to S6J). Taken together these results indicate that a reduction in PARIS restores the mitochondrial respiratory and biogenesis defects due to the PINK1 deficiency in human DA neurons without improving the mitophagy defects.

**Figure 7.**
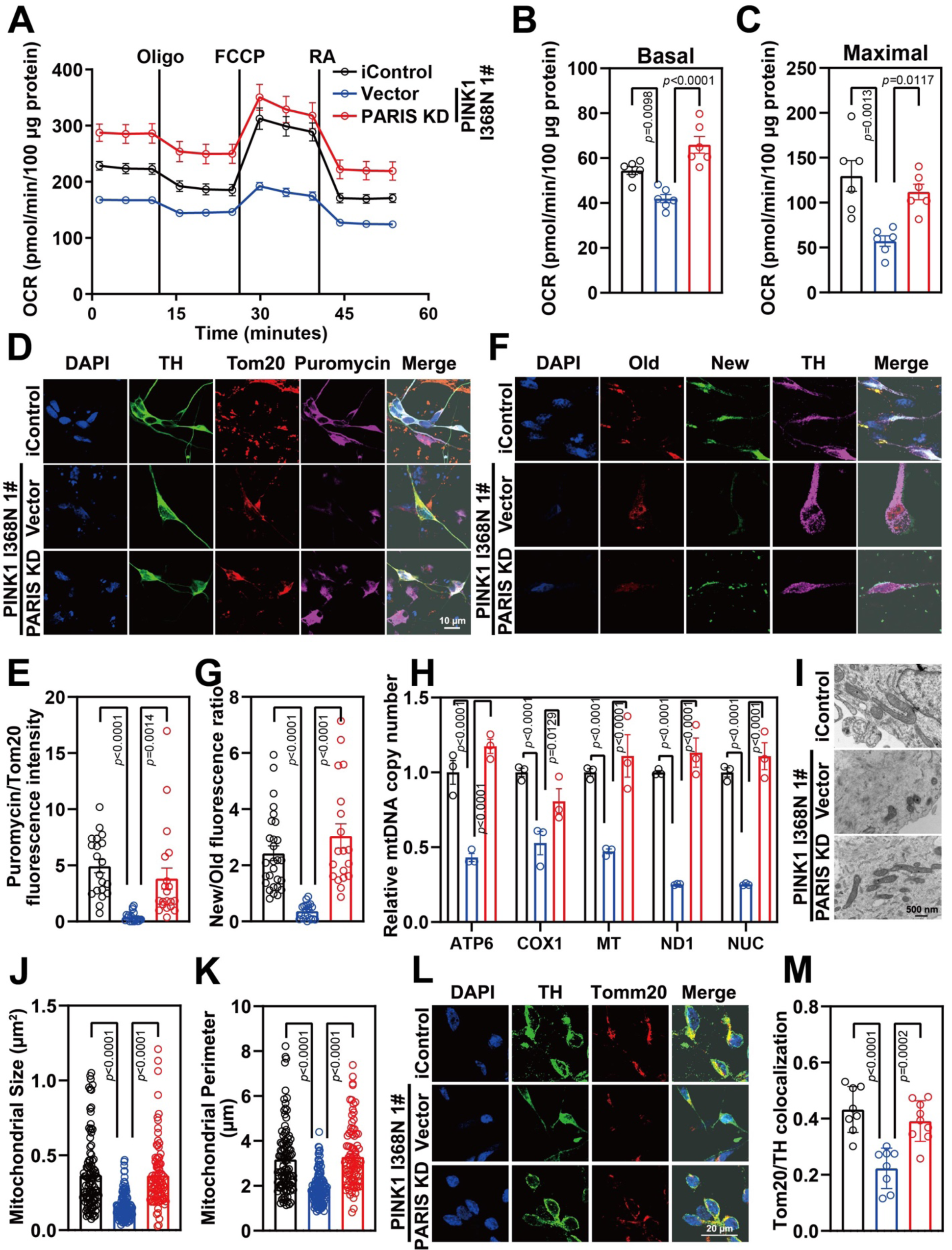
Mitochondrial function recused in PARIS knockdown in PINK1 p.I368N-1# mutant DA neurons. C. Mitochondrial oxygen consumption rate curve generated using Seahorse platform showing the mitochondrial function of isogenic control and PINK1 p.I368N 1# mutant neurons with or without PARIS knockdown. D. Quantification of basal respiration in isogenic control and PINK1 p.I368N 1# mutant neurons with or without PARIS knockdown. N=3 independent experiments. E. Quantification of maximal respiration in isogenic control and PINK1 p.I368N 1# mutant neurons with or without PARIS knockdown. N=3 independent experiments. F. Representative confocal images of puromycin labelling SUnSET assay in isogenic control and PINK1 p.I368N 1# mutant DA neurons with or without PARIS knockdown. G. Immunofluorescence intensity quantification of Tom20 and puromycin within TH positive neurons with or without PARIS knockdown. N=20 TH positive neurons in each group. H. Representative confocal images of SNAP-TAG Cox8a labeled mitochondria with TH staining in isogenic control and PINK1 p.I368N 1# mutant neurons with or without PARIS knockdown. I. Fluorescence intensity quantification of old and new mitochondria labeled by Cox8a in TH positive neurons with or without PARIS knockdown. N=20 TH positive neurons in each group. J. Relative mitochondrial DNA copy number quantification of ATP6, NUC, ND1, MT, and COX1 from isogenic control and PINK1 p.I368N-1# mutant neurons with or without PARIS knockdown. N=3 independent experiments. K. Representative transmission electron microscopy images of neurons from isogenic control and PINK1 p.I368N 1# mutant neurons with or without PARIS knockdown. N=15-30 cells in each group. L. Quantification of mitochondrial size per cell body from isogenic control and PINK1 p.I368N 1# mutant neurons with or without PARIS knockdown. N=15-30 cells in each group. M. Quantification of mitochondrial perimeter per cell body from isogenic control and PINK1 p.I368N-1# mutant neurons with or without PARIS knockdown. N. Representative confocal images of immunofluorescence staining of Tom20 and TH from isogenic control and PINK1 p.I368N 1# mutant neurons with or without PARIS knockdown. O. Quantification of colocalized Tom20 and TH in isogenic control and PINK1 p.I368N 1# mutant neurons with or without PARIS knockdown. N=10 TH positive neurons in each group.

## Discussion

The major finding of our study is that mitochondrial respiratory capacity and biogenesis reduction is due in large part to the PINK1 substrate, PARIS, elevation, and that neuronal mitochondrial dysfunction was unlikely due to mitophagy defects because the mitophagy defects were not reversed by reducing PARIS levels, while it restored the defects in mitochondria biogenesis and respiratory capacity.

Mitochondria are critical to cellular health and metabolism, and their dysfunction is amplified by defective quality control mechanisms and it is thought to play a major role in PD pathogenesis [10, 30]. In such mitochondrial quality control processes (that is, proteostasis, biogenesis, dynamics and mitophagy), mitochondrial biogenesis and mitophagy (degradation of damaged mitochondria) precisely regulates mitochondrial numbers [30]. PINK1/parkin are known to co-regulate mitochondrial mitophagy and biogenesis in mouse and Drosophila [31–34]. Consistent with these findings, we describe here that both mitophagy and mitochondria biogenesis defects were observed in human DA neurons lacking PINK1.

The use of human iPSCs derived DA neurons provided the ability to study the relative contribution of mitochondrial biogenesis versus mitophagy in the pathogenesis of PD resulting from loss of PINK1. PGC-1α protein levels were restored by reducing PARIS via CRISPR/Cas9, while mitophagy evaluated by using MitoKeima was not significantly reversed. Therefore, defects in mitophagy are not likely to be the major driver of mitochondrial dysfunction but mitochondrial biogenesis deficits are more likely occurring through PARIS upregulation and further downregulation of PGC-1α.

Compared with isogenic control DA neurons, the number of isogenic PINK1 mutant DA neurons were decreased, which was restored by downregulation of PARIS levels. These human DA neuron data are consistent with prior findings that downregulation PARIS levels in adult conditional knockdown of PINK1 mice can rescue the defects in mitochondrial biogenesis and prevent DA neurons loss when PGC-1α are restored. We also observed that PARIS level are upregulated leading to impaired mitochondrial biogenesis, depolarization and respiration that were rescued by the reduction of PARIS while failing to rescue mitophagy defects.

We evaluated the mitophagy by using the MitoKeima reporter assay and Western blot analysis of mitophagy markers LC3A and p62. Our results indicate that there is defective mitophagy in human DA neurons, which strengthens the notion that PINK1/PARKIN pathway regulates mitophagy in DA neurons. The findings reported here are also consistent with the notion that PINK1/PARKIN regulate mitophagy in immortalized cell lines, *Drosophila*, and rodent neurons [11, 12]. Previous in vivo studies observed no effect on basal mitophagy in PINK1 or parkin deficient Drosophila despite DA neurons loss [35, 36]. However, by using MitoKeima imaging and correlative light and electron microscopy (CLEM), basal mitophagy reduction was observed in 3 and 4 weeks old PINK1 or parkin deficient Drosophila [37], but there were no mitophagy defects in 1-week-old flies lacking PINK1 [37] despite mitochondria abnormalities displayed at this age [38]. These findings suggest that the mitophagy defects are separate from the mitochondrial deficits, which is similar to human parkin [22] and PINK1 deficient DA neurons. Interestingly, the PARIS homologue [39] in Drosophila also can drive the mitochondrial defects like it does within mouse and human DA neurons [18, 40].

IF mitophagy defects were the driver of mitochondrial deficits in either PINK1 or parkin deficient Drosophila and mice, mitochondrial mass would be observed to increase due to inefficient damaged mitochondria clearance. In contrast, in PINK1 and PARKIN mutant files [41], in PINK knockout mice [42] mitochondrial do not accumulate. Moreover, by crossing germline PARKIN knockout mice with PolgA^D257A^ mitochondrial mutator mouse, DA neurodegeneration and mitochondrial dysfunction did not seem to be due to mitophagy defects since there was no increase in mitochondrial mass [43]. Mitochondrial dysfunction and dopaminergic neurodegeneration with corresponding nigral-striatal neurobehavioral deficits were not observed in a similar study, supporting the notion of lacking synergism of mitochondrial dysfunction in mouse models of mitochondrial deficits [44]. Similarly, mitochondrial mass or mitophagy was not observed in germline parkin knockout mice crossed to the PD-mito-PstI mouse [45] or TFAM knockout mouse [46]. Taken together, these findings suggest that mitophagy impairment by loss of PINK1/PARKIN is not driving the alternations in mitochondrial function ultimately leading to DA neuron loss [11, 12]. On the other hand, by employing proteomic approaches, mitochondrial proteins reduction [47] and respiration defect [48] were observed in germline parkin knockout mice. Mitochondrial size, mass and number were decreased, which is consistent with mitochondrial biogenesis defects in human DA neurons lacking PINK1, while defects in mitochondrial biogenesis were restored but without restoration in mitophagy when PARIS levels were reduced. These results support our notion that PARIS levels elevation causes defects in mitochondrial biogenesis via downregulating PGC-1α, which is the driving force of the mitochondrial phenotype. Additionally, adult conditional knockout of parkin leads to reduced mitochondrial size, mass and number, and the reduction in mitochondrial mass can be prevented by knockdown of PARIS using shRNA [15]. Emerging evidence suggest that mitophagy can occur in PINK1/PARKIN-independent manner. At present, we cannot rule out the possibility that this PINK1/Parkin-independent mitophagy pathway plays a role in PD pathogenesis.

In conclusion, our results strengthen the idea that both mitochondrial mitophagy and biogenesis are impaired in PINK1 deficient human DA neurons, and that mitochondrial biogenesis impairment is the predominate driving force of mitochondrial function reduction and the ultimate loss of DA neurons in PD due to PINK1 deficiency. Therapeutic approaches to enhance mitochondrial biogenesis may offer a way to prevent DA neurons loss in PD.

### Author contributions

Conceptualization, H.W., V.L.D., T.M.D.; Methodology, H.W., R.C., L.X., M.K., J.A.-C.; Validation, H.W.; Formal Analysis and Investigation, H.W., R.C., L.X.; Resources, J.S., D.K., Z.K.W.,V.L.D., T.M.D.; Writing-Original Draft, H.W., V.L.D., T.M.D.; Writing-Review and Editing, H.W.,V.L.D., T.M.D.; Supervision, V.L.D., T.M.D.; Project Administration, V.L.D., T.M.D.; Funding Acquisition, V.L.D., T.M.D.

### Competing interests

The value of patents owned by Valted, LLC could be affected by the study described in this article. Also, Dr. T. Dawson and V. Dawson are founders of Valted, LLC and hold an ownership equity interest in the company. This arrangement has been reviewed and approved by the Johns Hopkins University in accordance with its conflict of interest policies. ZKW serves as PI or Co-PI on Biohaven Pharmaceuticals, Inc. (BHV4157-206) and Vigil Neuroscience, Inc. (VGL101-01.002, VGL101-01.201, PET tracer development protocol, Csf1r biomarker and repository project, and ultra-high field MRI in the diagnosis and management of CSF1R-related adult-onset leukoencephalopathy with axonal spheroids and pigmented glia) projects/grants. He serves as Co-PI of the Mayo Clinic APDA Center for Advanced Research and as an external advisory board member for the Vigil Neuroscience, Inc., and as a consultant on neurodegenerative medical research for Eli Lilli & Company.

## Acknowledgments

The authors thank Noelle Burgess for creating and assisting with illustrations. We thank the Dawson lab personnel for helpful suggestions and Weiran Chen and Ahmet Hoke for access to a Seahorse instrument.This work was supported by grants from JPB Foundation. The authors acknowledge the joint participation by the Adrienne Helis Malvin Medical Research Foundation and the Diana Helis Henry through its direct engagement in the continuous active conduct of medical research in conjunction with The Johns Hopkins Hospital and the Johns Hopkins University School of Medicine and the Foundation’s Parkinson’s Disease Programs M-1, M-2, H-2014, M-2016. T.M.D. is the Leonard and Madlyn Abramson Professor in Neurodegenerative Diseases. ZKW is partially supported by the NIH/NIA and NIH/NINDS (1U19AG063911, FAIN: U19AG063911), Mayo Clinic Center for Regenerative Medicine, the gifts from the Donald G. and Jodi P. Heeringa Family, the Haworth Family Professorship in Neurodegenerative Diseases fund, and The Albertson Parkinson’s Research Foundation.

## Methods & Materials

### hiPSC maintenance and DA differentiation

Human iPSCs isogenic control (NN0005032) and PINK1 p.I368N-1# mutant (NN0005033), SC1014 and SC1015 (wildtype control), PINK1 p.Q456X (1046#), and PINK1 p.I368N-2# (1067#) were cultured using mTeSR (STEMCELL Technologies, Catlog #85850) media on Matrigel (Corning, Catlog #354277) coated 12-well plate. DA differentiation protocol was optimized as previously published. Briefly, hiPSCs were dissociated with Collagenase Type IV (STEMCELL Technologies, Catlog #07909). Differentiation media contain growth factors and small molecule including SHH (100 ng/ml, PeproTech, Catalog #100-45), FGF8b (100 ng/ml, PeproTech, Catalog #100-25), SB431542 (10 µM, Stemgent, Catlog #04-0010-05), LDN193189 (100 nM, Stemgent, Catlog #04-0074-02), CHIR99021 (3 µM, Stemgent, Catlog #04-0004-02) and Purmorphine (2 µM, Stmgent, Catalog #04-0009); and mature medium containing BDNF (20 ng/ml, PeproTech, Catalog #450-02), GDNF (20 ng/ml, PeproTech, Catlog #450-10), TGF-β3 (1 ng/ml, PeproTech, Catlog #100-36E), ascorbic acid (200 µM, Sigma-Aldrich, Catlog #A4034), cAMP (500 µM, Sigma-Aldrich, Catalog #D0627) and DAPT (10 µM, Cell Signaling Technology, Catlog # 15020S) were changed every other day.

### Immunofluorescence

NPCs were seeded onto polyornithine and laminin (100 µg/ml) precoated glass coverslips in 24 well plate, then started to differentiate using conditioned differentiation medium maturation around day 60. After treatment or labelling, cells were fixed with 4% (w/v) paraformaldehyde for 15 mins, washed with PBS and permeabilized with 0.5% Triton X-100 for 15 mins. After blocking with 5% donkey serum plus 5% bovine serum albumin (BSA) in PBS for 1 h, fixed cells were incubated with primary antibodies (1:500 dilution) in blocking buffer overnight at 4 °C. Following 3 times PBS washing for 5 min each, samples were incubated with fluorochrome-conjugated secondary antibody (donkey anti-rabbit Alexa Fluor-488/Alexa Fluor-568/Alexa Fluor-647, anti-mouse Alexa Fluor-568/Alexa Fluor-568/Alexa Fluor-647, 1:500-1:1000 dilution) plus DAPI for 1-2 h in dark at 37 °C incubator. Rinse coverslips 3 times in PBS for each 5 min each and finally mounted onto microscope slide. High resolution images were captured with ZEISS LSM 880 confocal microscope.

### Protein extraction, SDS-PAGE and Western blot

Cells were washed with PBS and collected using scrapper, soluble and insoluble fractions of cell pellet were prepared by homogenization with lysis buffer containing 1% Triton X-100 with phosphatase inhibitor cocktail and protease inhibitor mixture in PBS. Cell lysates were centrifuged at 12000 rpm for 30 mins at 4 °C. The resulting supernatant was removed and transferred to a fresh Eppendorf tube as soluble fraction. The resulting pellet was washed with lysis buffer one time and homogenized and sonicated with lysis buffer containing 1% SDS. After centrifugation at 4 °C, 12000 rpm for 20 mins, the resulting supernatant was carefully removed as insoluble fraction. The protein concentration was determined by the Pierce BCA analysis. Protein lysis with Laemmli sample buffer were separated by 4-20% polyacrylamide gels and transferred onto polyvinylidene difluoride (PVDF) membranes. The membranes were blocked with 5% non-fat milk in PBST and then incubated with primary antibodies against the PARIS (Proteintech, #24542-1-AP, 1:2500), PGC-1α (Novus, #NBP1-04676, 1: 2000), TH (Millipore, #AB152, 1:2000), TUJ1 (Biolegend, #801201, 1:3000), PINK1 (Novus, #NBP2-36488, 1:2000), PARKIN (Cell Signaling, #4211, 1:2000), LC3A (Cell Signaling, #4599, 1:2000), P62 (Cell Signaling, #8025, 1:1000), CytC (Cell Signaling, #4280, 1:1000), CoxIV (Cell Signaling, #4850, 1:1000) and β-actin (Sigma, #A-11029, 1:3000). After washing with PBST 3 times for each 5 min, the following blots were incubated with peroxidase-conjugated goat anti-rabbit or mouse IgG (Thermofisher, #A-11036 or #A-11029, 1: 3000) for 1 h at room temperature. Blots were visualized on Amersham Imager 600 by chemiluminescence method. The signal intensity of the band was quantified by Image J software.

### Sea Horse assay

Seahorse assay was performed using the Seahorse XF96 analyzer (Agilent) and the XF Cell Mito Stress Test Kit according to the manufacturer’s instructions. Briefly, NPCs (50000 cells per well) were seeded to polyornithine and laminin (100 µg/ml) precoated 96-well Seahorse plate, and further differentiated for at least 14 days for further mature. Before the start of assay, Agilent Seahorse XFe96 cartridge (Agilent, 102416-100) was hydrated with 200 μl of calibrant solution (Agilent, 100840-000) overnight at 37 °C. The next day, neurons were washed twice with assay buffer containing with glucose (5 mM), pyruvate (2 mM) and glutamine (2 mM). After the second wash, neurons were incubated with assay media and further incubated at 37 °C in a non-CO_2_ incubator to equilibrate 30 mins. Oxygen consumption rate (OCR) was measured under basal conditions, and the sequential injection of 2 µM Oligomycin, 2 µM FCCP and 0.5 µM Rotenone. The instrument for Mito Stress Test procedures consists of 1 min mixing, 0 min wait and 3 min measurement. After OCR recording, the concentration of protein lysis from each well was measured and used to normalize OCR values of each well. OCR values were calculated by the Wave (Seahorse XF96 software), the results are expressed as mean ± standard errors of the mean (SEM).

### Mitochondrial DNA quantitative PCR

Total DNA was extracted from differentiated DA neurons using QIAGEN blood & tissue genomic DNA isolation kit following the protocol provided. Purified DNA concentration was measured using nanodrop spectrophotometer. For qPCR, 10 µl qPCR reaction volume contained 100 ng genomic DNA template, SYBR Green Mix, 0.3 µM primers of each and nuclease-free water. Thermal cycling was run on thermal cycler (Applied Biosystems) by first heating to 95 °C for 3 min, then 40 cycles of 95 °C 15 sec, 58 °C 30 sec and 72 °C 30 sec. Primers were listed in Table S1. All sample included triplicate reactions; averaged Ct values were used for further analysis. GAPDH was used as internal control.

### Mitochondrial membrane potential assay

Differentiated DA neurons mitochondrial membrane potential were determined with MitoTracker Red CMXRos following the manufacturer’s protocols. Briefly, DA neurons on coverslip were incubated DA media containing 75 nM tracker for 30 min at 37 °C incubator. Following 3 times washing with PBS, neurons were fixed with 4% PFA for 15 min and then further stained with primary and secondary antibodies following the protocol of immunofluorescence described above. Images were captured with ZEISS LSM 880 confocal microscope.

### Lentiviral transduction

The CRISPR/Cas9 based PARIS knockdown virus were packaged using lentiviral transfer plasmid constructed as our previous published paper. Matured DA neurons were infected with lentivirus encoding gRNA, and its knockdown efficiency was confirmed by western blot.

### SNAP-Tag assay

Specific mitochondrial labelling has been successfully conducted via fusion of mitochondrial protein Cox8A with SNAP. NPCs were seeded onto glass coverslips and differentiated around day 55. Lentiviral SNAP tag Cox8a was prepared and transduced to human DA neurons for another 3-4 days. After 3 times wash with media, neurons were incubated with SNAP-tag first substrate (TMR-Star) containing DA medium, cultured for another 1 h and then 3 times wash with media. After 24 h culture, the second substrate (Oregon Green) was added into well and maintained for 1 h. Finally, cells were washed with media 3 times and then fixed with 4% PFA and followed by further immunostaining processing.

### SUnSET assay

SUnSET assay was performed to examine the neuronal global protein synthesis via puromycin labeling. Incorporated puromycin in active nascent polypeptide chain can be detected using antibody. Human NPCs were seeded onto glass coverslips and then differentiated into matured DA neurons. Fresh puromycin (1 µM) containing DA culture media was added into each well and incubated for 1 at 37 °C incubator. After 3 times wash with PBS to remove unlabeled puromycin, the samples were fixed with 4% PFA for further immunostaining procedures.

### Transmission electron microscopy (TEM) assay

NPCs were seeded on polyornithine and laminin (100 µg/ml) precoated 6-well plate and differentiated to mature DA neurons with or without PARIS KD lentivirus infection. Cells were washed with PBS and fixed in 2.5% glutaraldehyde, 3 mM MgCl_2_ in 100 mM sodium cacodylate buffer (pH 7.2) at 4 °C for 24 h, followed by buffer wash and incubated with 0.8% potassium ferrocyanide reduced osmium tetroxide in 100 mM sodium cacodylate buffer (pH 7.2) for 1 h on ice. Cells were stained with 2% uranyl acetate overnight and dehydrated by ethanol in cold room. Then, the samples were infiltrated and embedded in Epon/Araldite mixture for polymerization about 48 h at 37 °C. Thin sections were cut using a diamond knife and mounted on EM grids, then further stained with 2% uranyl acetate in 50% methanol containing lead citrate. Finally, samples were observed under Philips CM120 at 80 kV, and images were captured with high resolution camera. TEM images were analyzed using Image J software to determine the size and perimeter of mitochondria on randomly selected views (at least 15 cells in each group were analyzed).

### MitoKeima assay

NPCs were seeded on polyornithine and laminin (100 µg/ml) precoated glass coverslips in 24-well plate and differentiated to mature DA neurons. MitoKeima encoding lentivirus was prepared and transduced to human DA neurons for another 3-4 days. After 3 times wash with media, the coverslip was picked up and put into the live cell imaging chamber filled with same media of neurons maintained. Images were captured using 458-nm and 543-nm laser sequentially scanning under ZEISS LSM 880 confocal microscope.

### Statistical analysis

All quantitative data was presented as mean ± standard error of the mean (SEM) from 3 different independent experiments. Statistical comparisons were conducted by using GraphPad Prism software 7. Significance was tested by unpaired Student’s *t*-test, one-way analysis of variance and two-way ANOVA by Turkey’s post-hoc test or Bonferroni post-hoc test respectively, the *p* values are indicated in the figures and *p* < 0.05 was considered as significant.

### Data and material availability

All biological resources, antibodies, cell lines and model organisms and tools are either available through commercial sources or the corresponding authors. The data sets generated during and/or analyzed during the current study are available from the corresponding author on reasonable request.

