## Supplemental Files for "Defects in Mitochondrial Biogenesis Drive Mitochondrial Alterations in PINK1-deficient Human Dopamine Neurons"

### Supplementary Figure and Figure Legends

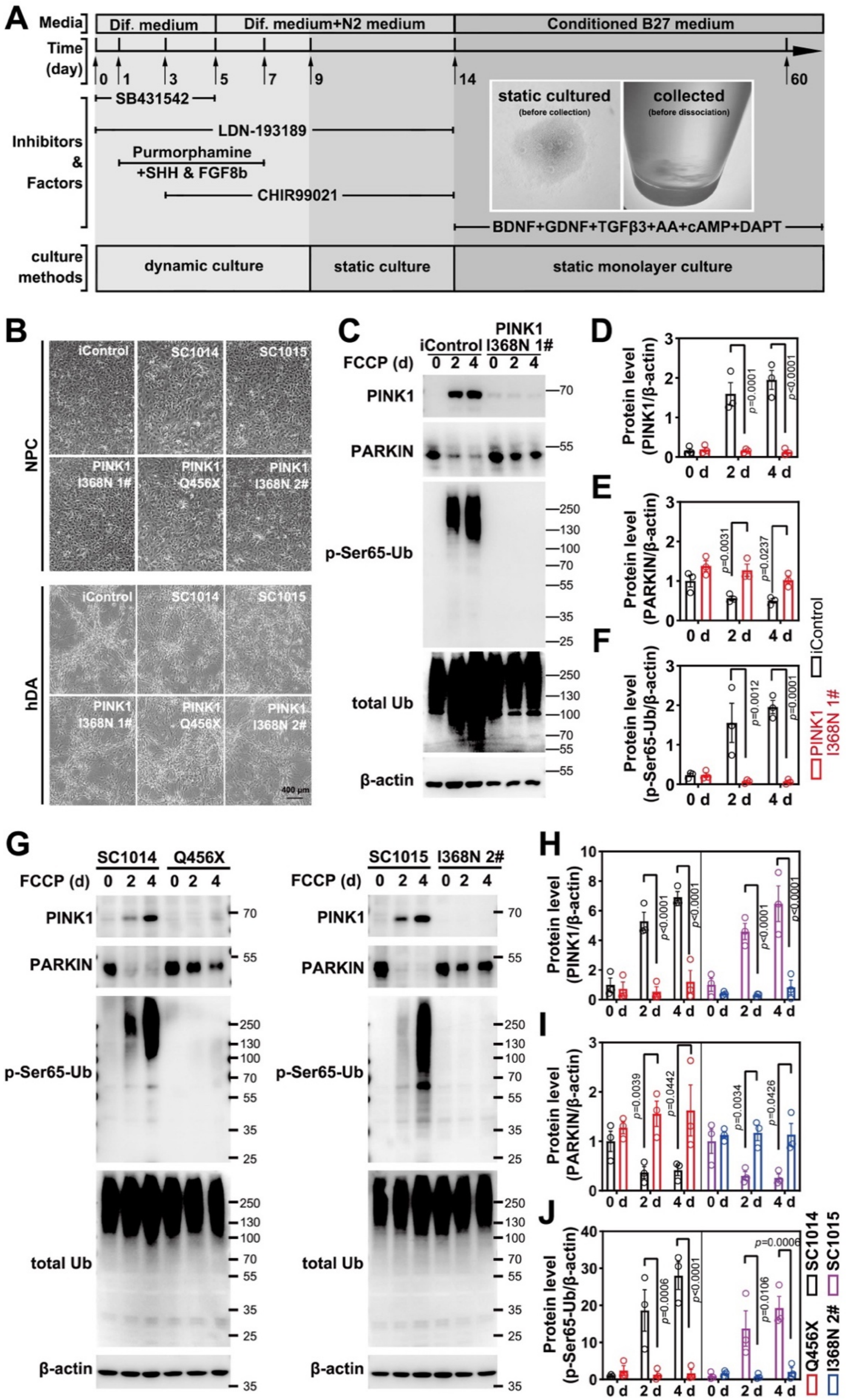

**Figure S1 Differentiation protocol optimization and DA neuron characterization**

A. Schematic illustration of hDA neuron differentiation protocol used, differentiation time, medium, factors or inhibitors and culture method were included, progenitor clumps were collected and dissociated before replating.
B. Representative images of progenitors and mature hDA neurons differentiated from isogenic control and PINK1 p.I368N 1# lines, wildtype SC1014 and SC1015, PINK1 p.Q456X and PINK1 p.I368N 2# mutant lines.
C. Immunoblot analysis of PINK1, PARKIN, pSer65-Ub, an Ub protein level in isogenic control and PINK1 p.I368N 1# neurons.
D. Quantification of PINK1 protein levels normalized by β-actin. N=3 independent experiments.
E. Quantification of PARIN protein levels by β-actin. N=3 independent experiments.
F. Quantification of pSer65-Ub protein levels normalized by β-actin. N=3 independent experiments.
G. Immunoblot analysis of PINK1, PARKIN, pSer65-Ub, an Ub protein level in wildtype SC1014 and SC1015, PINK1 p.Q456X and PINK1 p.I368N 2# neurons.
H. Quantification of PINK1 protein levels normalized by β-actin. N=3 independent experiments.
I. Quantification of PARIN protein levels by β-actin. N=3 independent experiments.
J. Quantification of pSer65-Ub protein levels by β-actin. N=3 independent experiments.

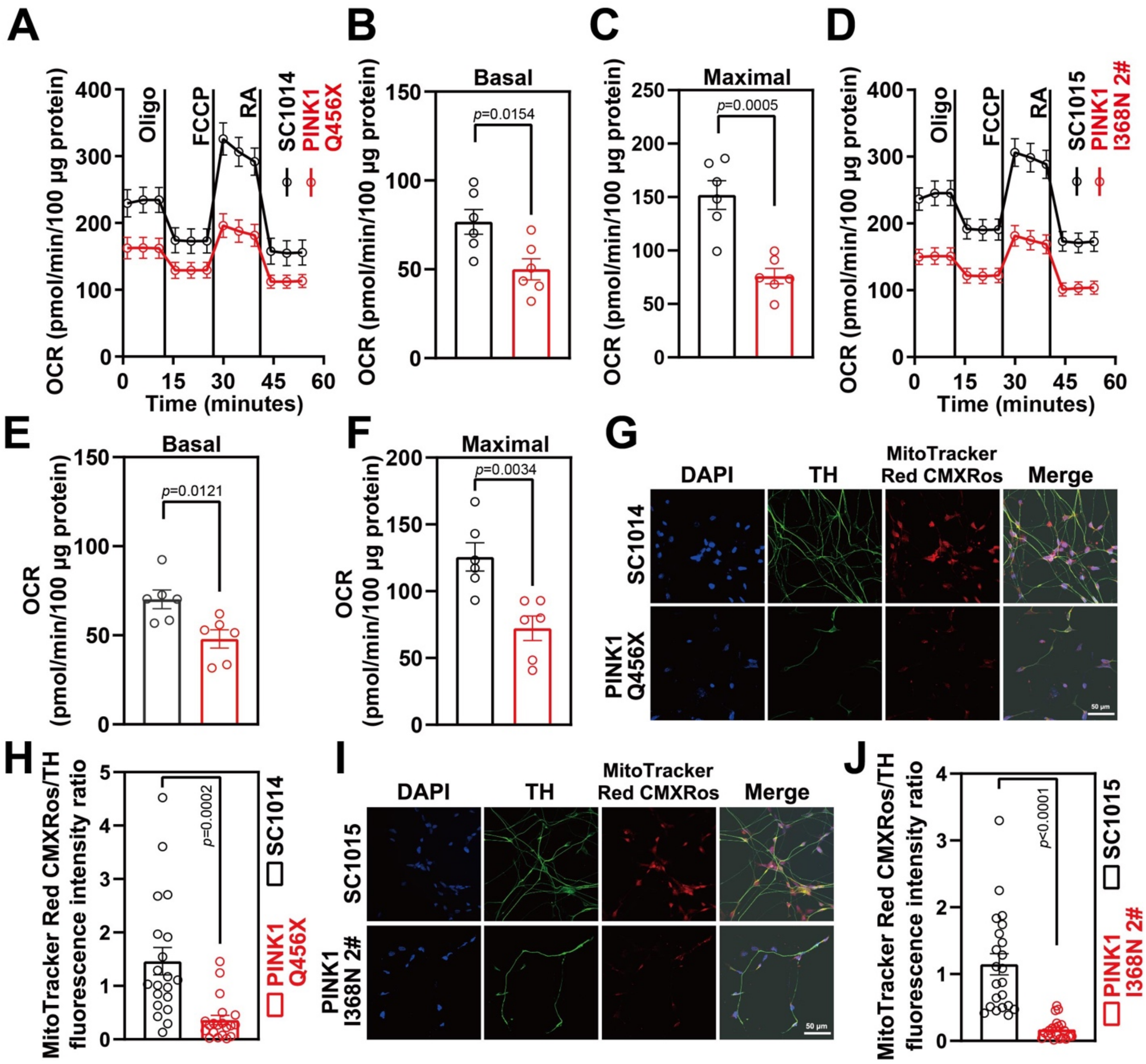


**Figure S2 Mitochondrial dysfunction in PINK1 p.Q456X mutant DA neurons**

A. Mitochondrial oxygen consumption rate curve generated using Seahorse platform showing the mitochondrial dysfunction in wildtype SC1014 and PINK1 p.Q456X mutant neurons in the presence of oligomycin, FCCP, and rotenone respectively.
B. Quantification of basal respiration in wildtype SC1014 and PINK1 p.Q456X mutant neurons. N=3 independent experiments.
C. Quantification of maximal respiration in wildtype SC1014 and PINK1 p.Q456X mutant neurons. N=3 independent experiments.
D. Mitochondrial oxygen consumption rate curve generated using Seahorse platform showing the mitochondrial dysfunction in wildtype SC1015 and PINK1 p.I368N 2# mutant neurons in the presence of oligomycin, FCCP, and rotenone respectively.
E. Quantification of basal respiration in wildtype SC1015 and PINK1 p.I368N-2# mutant neurons. N=3 independent experiments.
F. Quantification of maximal respiration in wildtype SC1015 and PINK1 p.I368N 2# mutant neurons. N=3 independent experiments.

G. Representative immunostaining images of TH-positive neurons and MitoTracker Red CMXRos from wildtype SC1014 and PINK1 p.Q456X mutant neurons.
H. Quantification of TH and MitoTracker Red CMXRos intensity. N=20 TH positive neurons in each group. N=15-30 TH positive neurons in each group.
I. Representative immunostaining images of TH-positive neurons and MitoTracker Red CMXRos from wildtype SC1015 and PINK1 p.I368N 2# mutant neurons.
J. Quantification of TH and MitoTracker Red CMXRos intensity. N=20 TH positive neurons in each group. N=15-30 TH positive neurons in each group.

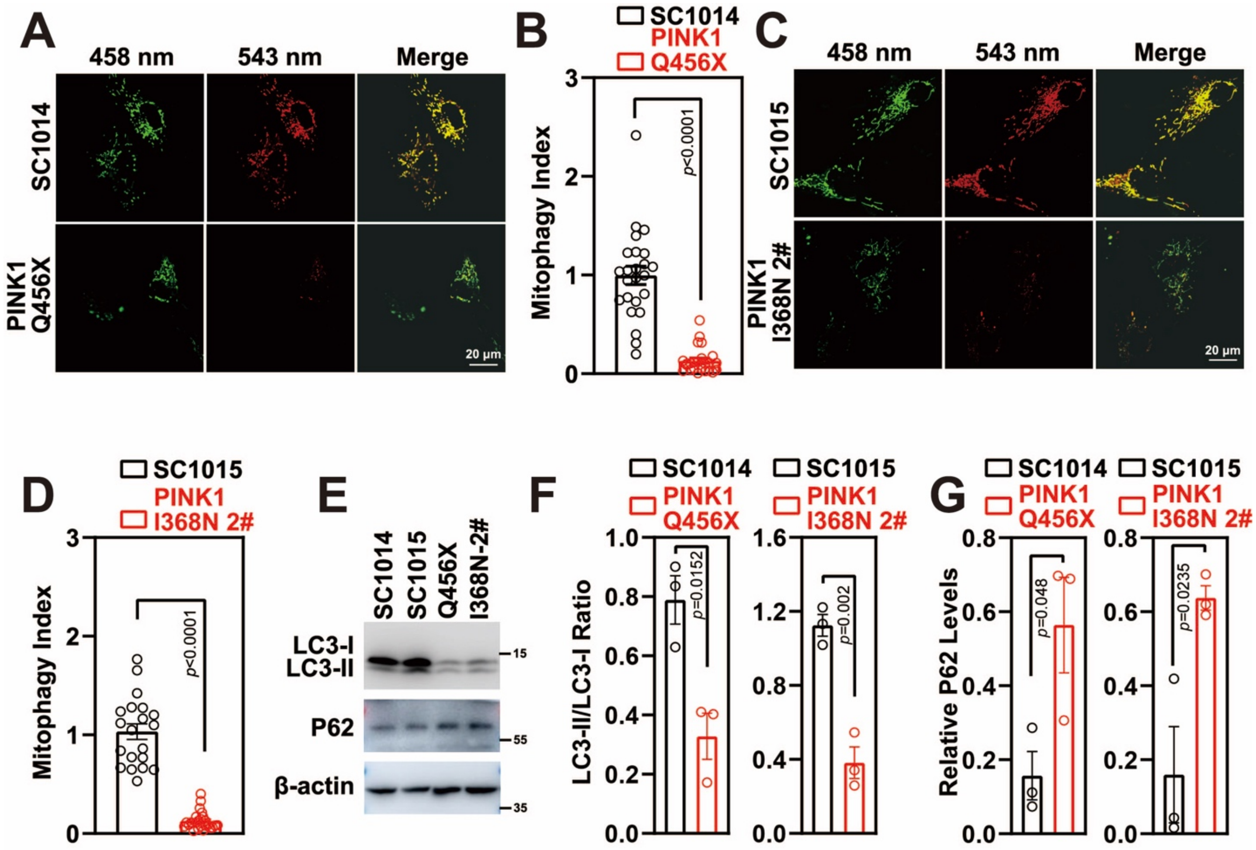


**Figure S3 Mitophagy defects in PINK1 p.Q456X mutant neurons**

A. Representative confocal live images of DA neurons infected with lentivirus encoding for MitoKeima.
B. Quantified mitophagy index in wildtype SC1014 and PINK1 p.Q456X mutant neurons. N=20 neurons in each group.
C. Representative confocal live images of DA neurons infected with lentivirus encoding for MitoKeima.
D. Quantified mitophagy index in wildtype SC1015 and PINK1 p.I368N 2# mutant neurons. N=20 neurons in each group.
E. Analysis of LC3-I/II and autophagy marker P62 protein level by immunoblot.
F. Quantification of immunoblot of LC3-I/II ratio are shown. N=3 independent experiments.
G. Quantification of immunoblot of P62 normalized to β-actin. N=3 independent experiments.

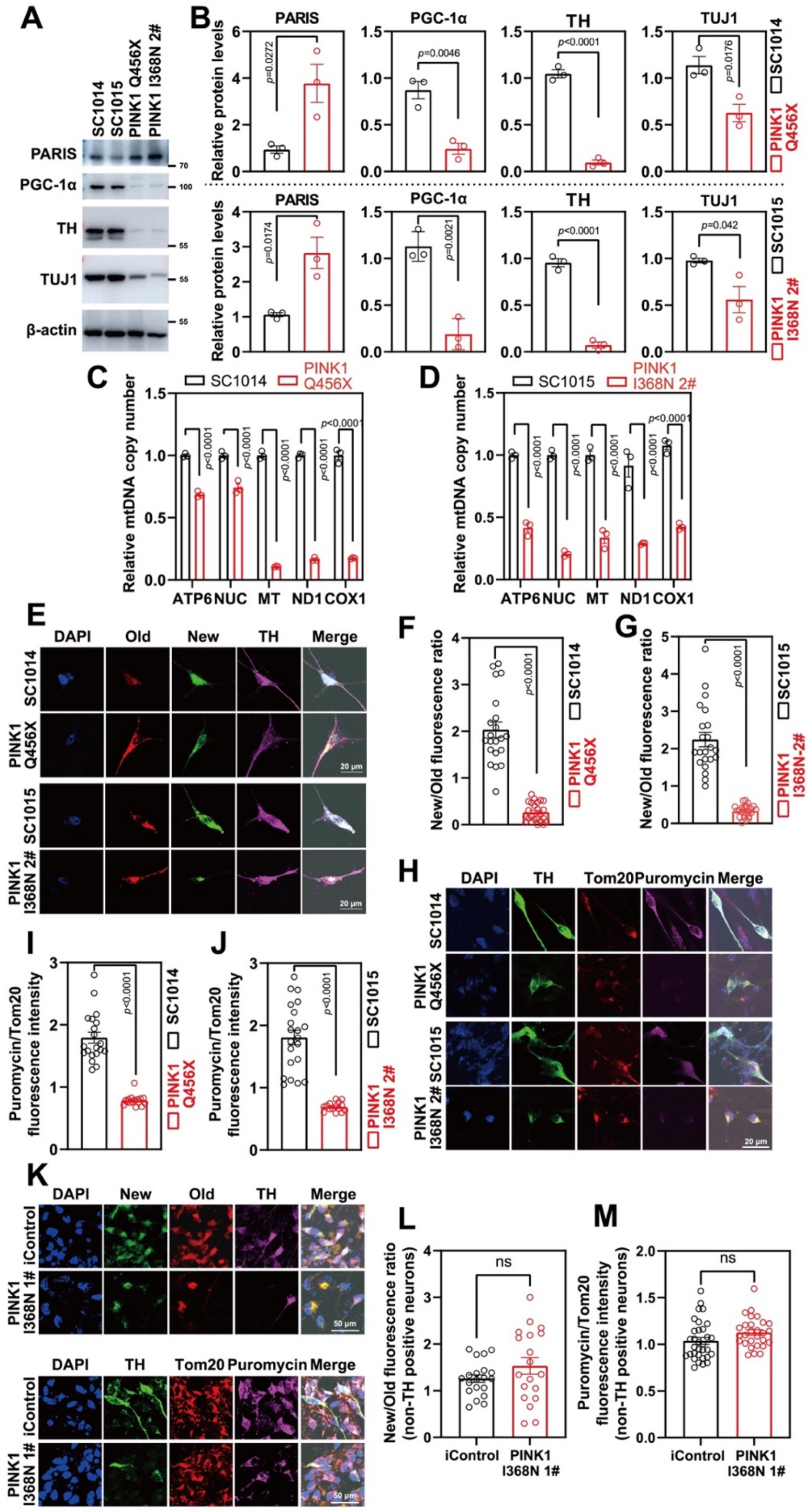


**Figure S4 Defects of mitochondrial biogenesis in PINK p.Q456X mutant DA neurons**

A. Analysis of PARIS, PGC-1α, TUJ1 and TH protein level in wildtype SC1014 and SC1015, PINK1 p.Q456X and PINK1 p.I368N 2# mutant neurons by immunoblot.
B. Quantification of immunoblot of PARIS, PGC-1α, TUJ1 and TH normalized to β-actin. N=3 independent experiments.
C. Representative confocal images of SNAP-TAG Cox8a labeled mitochondria with TH staining in wildtype SC1014 and SC1015, PINK1 p.Q456X and PINK1 p.I368N 2# mutant neurons.
D. Fluorescence intensity quantification of old and new mitochondria labeled by Cox8a in TH positive neurons. N=20 TH positive neurons in each group.
E. Fluorescence intensity quantification of old and new mitochondria labeled by Cox8a in TH positive neurons. N=20 TH positive neurons in each group.
F. Representative confocal images of puromycin labelling SUnSET assay in wildtype SC1014 and SC1015, PINK1 p.Q456X and PINK1 p.I368N 2# mutant neurons.
G. Immunofluorescence intensity quantification of Tom20 and puromycin within TH positive neurons. N=20 TH positive neurons in each group.
H. Immunofluorescence intensity quantification of Tom20 and puromycin within TH positive neurons. N=20 TH positive neurons in each group.
I. Representative confocal images of SNAP-TAG and SUnSET assay in isogenic control and PINK1 p.I368N 1# mutant neurons.
J. Fluorescence intensity quantification of old and new mitochondria labeled by Cox8a in non-TH positive neurons. N=20 TH positive neurons in each group.
K. Immunofluorescence intensity quantification of Tom20 and puromycin within non-TH positive neurons. N=20 TH positive neurons in each group.
L. Relative mitochondrial DNA copy number quantification of ATP6, NUC, ND1, MT, and COX1 in wildtype SC1014 and PINK1 p.Q456X mutant neurons. N=3 independent experiments.
M. Relative mitochondrial DNA copy number quantification of ATP6, NUC, ND1, MT, and COX1 in wildtype SC1015 and PINK1 p.I368N 2# mutant neurons. N=3 independent experiments.

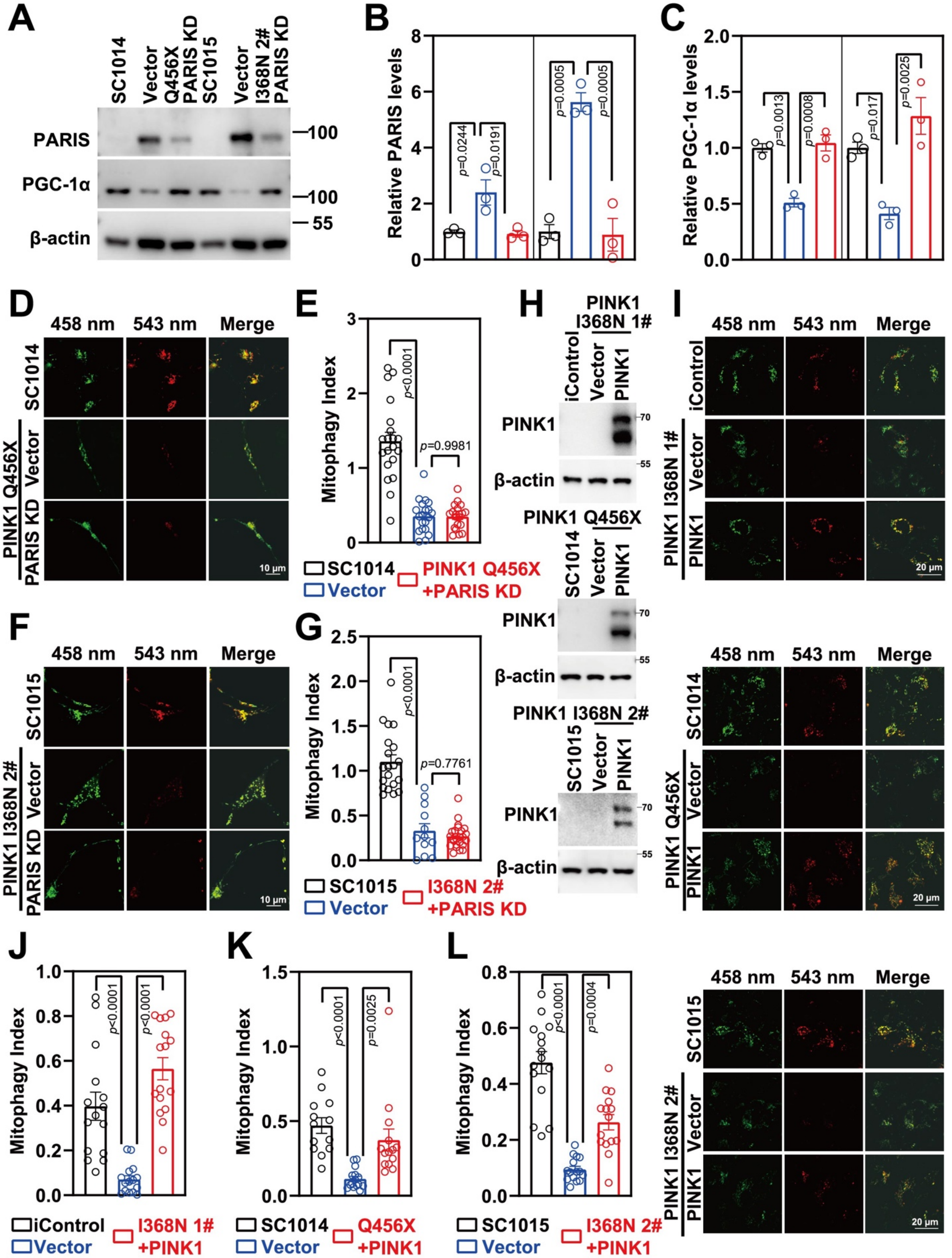


**Figure S5 Characterization of PARIS knockdown in PINK1 p.Q456X mutant neurons.**

A. Immunoblot analysis of PARIS, and PGC-1α protein level in wildtype SC1014 and SC1015, PINK1 p.Q456X and PINK1 p.I368N 2# mutant neurons with or without PARIS knockdown.
B. Quantified intensities of PARIS are shown, relative protein level was normalized to β-actin. N=3 independent experiments.
C. Quantified intensities of PGC-1α are shown, relative protein level was normalized to β-actin. N=3 independent experiments.
D. Representative confocal images of wildtype SC1014 and PINK1 p.Q456X mutant neurons infected with lentivirus encoding Mito-Keima mitophagy reporter systems.
E. Quantified mitophagy index in wildtype SC1014 and PINK1 p.Q456X mutant neurons with or without PARIS knockdown. N=20 TH positive neurons in each group.
F. Representative confocal images of wildtype SC1015 and PINK1 p.I368N 2# mutant neurons infected with lentivirus encoding Mito-Keima mitophagy reporter systems.
G. Quantified mitophagy index in wildtype SC1015 and PINK1 p.I368N-2# mutant neurons with or without PARIS knockdown. N=20 TH positive neurons in each group.
H. Immunoblot analysis of PINK1 protein level in isogenic control and PINK1 p.I368N 1# mutant, wildtype SC1014 and SC1015, PINK1 p.Q456X and PINK1 p.I368N-2# mutant neurons with or without PINK overexpression.
I. Representative confocal images of isogenic control and PINK1 p.I368N 1# mutant, wildtype SC1014 and SC1015, PINK1 p.Q456X and PINK1 p.I368N-2# mutant neurons infected with lentivirus encoding Mito-Keima mitophagy reporter systems.
J. Quantified mitophagy index in isogenic control and PINK1 p.I368N 1# mutant neurons with or without PINK overexpression. N=20 TH positive neurons in each group.
K. Quantified mitophagy index in wildtype SC1014 and PINK1 p.Q456X mutant neurons with or without PINK overexpression. N=20 TH positive neurons in each group.
L. Quantified mitophagy index in wildtype SC1015 and PINK1 p.I368N 2# mutant neurons with or without PINK overexpression. N=20 TH positive neurons in each group.

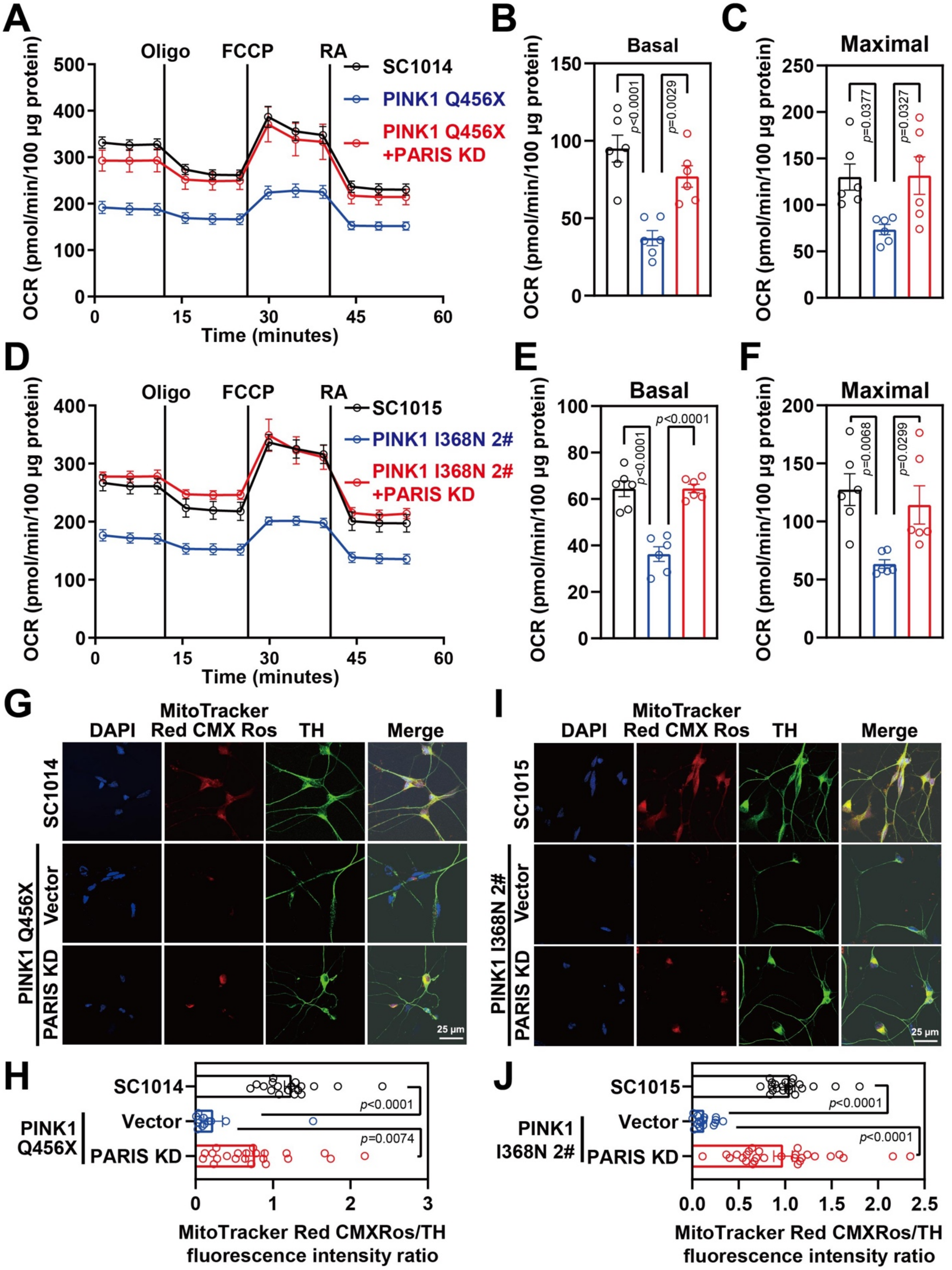


**Figure S6 Mitochondrial function recused in PINK1 p.Q456X mutant neurons by PARIS**

**knockdown**

A. Mitochondrial oxygen consumption rate curve generated using Seahorse platform showing the mitochondrial function of wildtype SC1014 and PINK1 p.Q456X mutant with or without PARIS knockdown.
B. Quantification of basal respiration in wildtype SC1014 and PINK1 p.Q456X mutant neurons with or without PARIS knockdown. N=3 independent experiments.
C. Quantification of maximal respiration in wildtype SC1014 and PINK1 p.Q456X mutant neurons with or without PARIS knockdown. N=3 independent experiments.
D. Mitochondrial oxygen consumption rate curve generated using Seahorse platform showing the mitochondrial function of wildtype SC1015 and PINK1 p.I368N 2# mutant neurons with or without PARIS knockdown.
E. Quantification of basal respiration in wildtype SC1015 and PINK1 p.I368N 2# mutant neurons with or without PARIS knockdown. N=3 independent experiments.
F. Quantification of maximal respiration in wildtype SC1015 and PINK1 p.I368N 2# mutant neurons with or without PARIS knockdown. N=3 independent experiments.
G. Representative immunostaining images of TH-positive neurons and MitoTracker Red CMXRos from wildtype SC1014 and PINK1 p.Q456X mutant neurons with or without PARIS knockdown.
H. Quantification of TH and MitoTracker Red CMXRos intensity. N=20 TH positive neurons in each group. N=15-30 TH positive neurons in each group.
I. Representative immunostaining images of TH-positive neurons and MitoTracker Red CMXRos from wildtype SC1015 and PINK1 p.I368N 2# mutant neurons with or without PARIS knockdown.
J. Quantification of TH and MitoTracker Red CMXRos intensity. N=20 TH positive neurons in each group. N=15-30 TH positive neurons in each group.

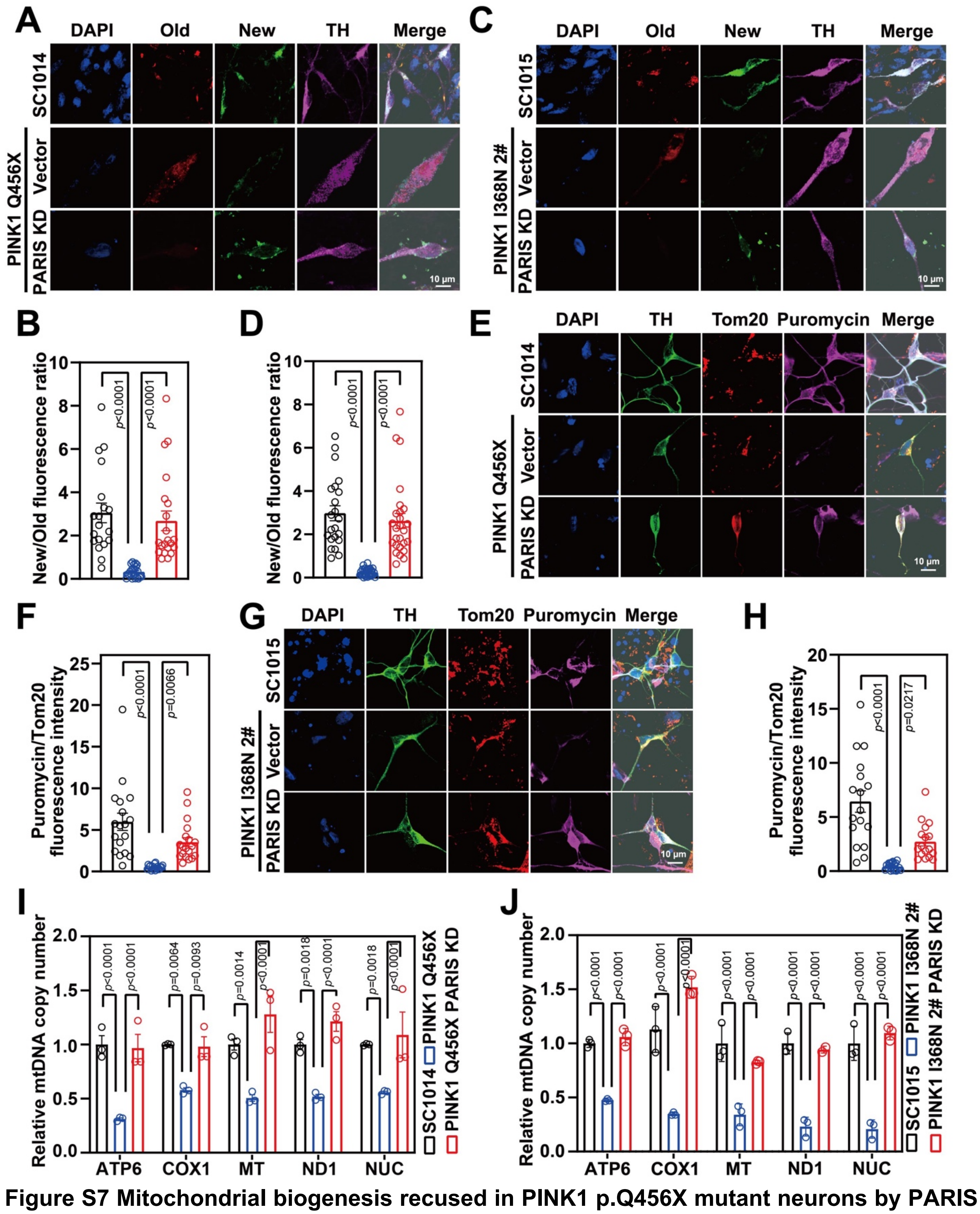


**Figure S7 Mitochondrial biogenesis recused in PINK1 p.Q456X mutant neurons by PARIS knockdown**

A. Representative confocal images of SNAP-TAG Cox8a labeled mitochondria with TH staining in wildtype SC1014 and PINK1 p.Q456X mutant neurons.

B. Fluorescence intensity quantification of old and new mitochondria labeled by Cox8a in TH positive neurons. N=20 TH positive neurons in each group.
C. Representative confocal images of SNAP-TAG Cox8a labeled mitochondria with TH staining in wildtype SC1015 and PINK1 p.I368N 2# mutant neurons.
D. Fluorescence intensity quantification of old and new mitochondria labeled by Cox8a in TH positive neurons. N=20 TH positive neurons in each group.
E. Representative confocal images of puromycin labelling SUnSET assay in wildtype SC1014 and PINK1 p.Q456X mutant neurons with or without PARIS knockdown.
F. Immunofluorescence intensity quantification of Tom20 and puromycin within TH positive neurons with or without PARIS knockdown. N=20 TH positive neurons in each group.
G. Representative confocal images of puromycin labelling SUnSET assay in wildtype SC1015 and PINK1 p.I368N 2# mutant neurons with or without PARIS knockdown.
H. Immunofluorescence intensity quantification of Tom20 and puromycin within TH positive neurons with or without PARIS knockdown. N=20 TH positive neurons in each group.
I. Relative mitochondrial DNA copy number quantification of ATP6, NUC, ND1, MT, and COX1 from wildtype SC1014 and PINK1 p.Q456X mutant neurons with or without PARIS knockdown. N=3 independent experiments.
J. Relative mitochondrial DNA copy number quantification of ATP6, NUC, ND1, MT, and COX1 from wildtype SC1015 and PINK1 p.I368N 2# mutant neurons with or without PARIS knockdown. N=3 independent experiments.

**Supplementary Table**

**Table S1. Primers used for RT-PCR**

| Gene | Sequence |
|---|---|
| ATP6 | F: CGCCACCCTAGCAATATCA |
| | R: TTAAGGCGACAGCGATTTC |
| MT | F: CCCCACAAACCCCATTACTAAACCCA |
| | R: TTTCATCATGCGGAGATGTTGGATGG |
| COX1 | F: CGATGCATACACCACATGA |
| | R: AGCGAAGGCTTCTCAAATC |
| NUC | F: ATACCCCCGATTCCGCTACGA |
| | R: GTTTGAGGGGGAATGCTGGAG |
| ND1 | F: ATACCCCCGATTCCGCTACGA |
| | R: GTTTGAGGGGGAATGCTGGAG |
| B2-micoglobulin | F: TGCTGTCTCCATGTTTGATGTATCT |
| | R: TCTCTGCTCCCCACCTCTAAGT |
